# Protein Design and Variant Prediction Using Autoregressive Generative Models

**DOI:** 10.1101/757252

**Authors:** Jung-Eun Shin, Adam J. Riesselman, Aaron W. Kollasch, Conor McMahon, Elana Simon, Chris Sander, Aashish Manglik, Andrew C. Kruse, Debora S. Marks

**Affiliations:** Department of Systems Biology, Harvard Medical School; Currently at insitro; Department of Biological Chemistry and Molecular Pharmacology, Harvard Medical School; Harvard College; Currently at Reverie Labs; Department of Cell Biology, Harvard Medical School and Department of Data Sciences, Dana Farber Cancer Institute; Department of Pharmaceutical Chemistry, University of California San Francisco; Department of Anesthesia and Perioperative Care, University of California San Francisco; Broad Institute of Harvard and MIT

## Abstract

The ability to design functional sequences and predict effects of variation is central to protein engineering and biotherapeutics. State-of-art computational methods rely on models that leverage evolutionary information but are inadequate for important applications where multiple sequence alignments are not robust. Such applications include the prediction of variant effects of indels, disordered proteins, and the design of proteins such as antibodies due to the highly variable complementarity determining regions. We introduce a deep generative model adapted from natural language processing for prediction and design of diverse functional sequences without the need for alignments. The model performs state-of-art prediction of missense and indel effects and we successfully design and test a diverse 10^5^-nanobody library that shows better expression than a 1000-fold larger synthetic library. Our results demonstrate the power of the ‘alignment-free’ autoregressive model in generalizing to regions of sequence space traditionally considered beyond the reach of prediction and design.

## Introduction

Over the past twenty years, success in protein engineering has emerged from two distinct approaches, directed evolution^1,2^ and knowledge-based force-field modeling^3,4^. Designing and generating biomolecules with known function is now a major goal of biotechnology and biomedicine, propelled by our ability to synthesize and sequence DNA at increasingly low costs. However, since the space of possible protein sequences is so large (for a protein of length 100 this is 10^130^), deep mutational scans^5^ and even very large libraries (e.g. >10^10^ variants) barely scratch the surface of the possibilities. As the vast majority of possible sequences will be non-functional proteins, it is crucial to minimize or eliminate these sequences from libraries. Therefore, the open challenge is to develop computational methods that can accelerate this search and bias the search space for protein sequences that are likely to be functional. This will enable design of libraries for tractable high-throughput experiments that are optimized for functional sequences and variants that are distant in sequence.

Antibody design is a particularly challenging problem in the area of statistical modeling of sequences for the purposes of prediction and design. Antibodies are valuable tools for molecular biology and therapeutics because they can detect low concentrations of target antigens with high sensitivity and specificity^6^. Single-domain antibodies, or nanobodies, are composed solely of the variable domain of the canonical antibody heavy chain. The increasing demand for and success with rapid and efficient discovery of novel nanobodies using phage and yeast display methods^7-10^ have spurred interest in the design of optimal starting libraries. Previous statistical and structural modeling of antibody repertoires^11-18^ have addressed the characterization of sequences of natural antibodies or predicted higher affinity sequences from immunization or selection experiments. One of the biggest challenges is to design libraries diverse enough to target many antigens but also be well-expressed, stable, and non poly-reactive. In fact, a large, state-of-art synthetic library contains a substantial fraction of non-functional proteins^8^ because library construction methods lack higher-order sequence constraints. Eliminating these non-functional proteins requires multiple rounds of selection and poses the single highest barrier to identifying high-affinity antibodies. In order to circumvent these limitations, there has been emphasis on very large libraries (∼10^9-1010^) to achieve these desired features^19, 20^.

Instead of experimentally producing unnecessarily massive, largely non-functional libraries, we can design smart libraries of fit and diverse nanobodies for the development of highly specific and possibly therapeutic nanobodies. One way to approach this is to leverage the information in natural sequences to learn constraints on specific amino acids in individual positions in a way that captures their dependency on amino acids in other positions. The sequences of these variants contain rich information about what contributes to a stable, functional protein, and in recent years generative models of these natural protein sequences have been powerful tools for the prediction of the first 3D fold from sequences alone^21, 22^, to generally more 3D structures and conformational plasticity^23, 24^, protein interactions^25-28^, and most recently, mutation effects^29-34^. However, these state-of-art methods and established methods^35-38^ rely on sequence families and alignments, and alignment-based methods are inherently unsuitable for the statistical description of the variable length, hypermutated complementarity determining regions (CDRs) of antibody sequences, which encode the diverse specific of binding to antigens. While antibody numbering schemes such as IMGT provide consistent alignments of framework residues, alignments of the CDRs rely on symmetrical deletions^39^. Alignment-based models are also unreliable for low-complexity or disordered proteins^40^ and cannot handle variants that are insertions and deletions. Indels make up 15-21% of human polymorphisms^41-43^, 44% of human proteins contain disordered regions longer than 30 amino acids^40, 44^, and both are enriched in association with human diseases such as cystic fibrosis, many cancers^45, 46^, cardiovascular and neurodegenerative diseases, and diabetes^47, 48^.

By contrast, the deep models that have transformed our ability to generate realistic speech such as text-to-speech^49, 50^ and translation^51, 52^ use generative models that do not require “word alignment”, e.g., between equisemantic sentences, but instead employ an autoregressive likelihood to tackle context-dependent language prediction and generation. Using this process, an audio clip is decomposed into discrete time steps, a sentence into words, and a protein sequence into amino acid residues. Models that decompose high-dimensional data into a series of steps predicted sequentially are termed autoregressive models, and they are well suited to variable-length data that have not been forced into a defined structure such as a multiple-sequence alignment. Autoregressive generative models are uniquely suited for modeling and designing the complex, highly diverse CDRs of antibodies. Here, we develop and apply a new autoregressive generative model that aims to capture key statistical properties of sets of sequences of variable lengths.

We first test our method on the problem of prediction of mutation effects, which are typically analyzed using alignment based statistical methods. The new method performs on par with the DeepSequence machine-learning VAE-based method^30^, which does require aligned sequences and which in an independent evaluation, testing against experimental data, was reported to outperform all currently available methods^34^. In addition to this state-of-the-art performance, our new alignment-free method is inherently more general. It can deal with a much larger class of sequences and take into account variable length effects. Another recently developed method^53^ does aim to quantify the of mutation effects without the need for alignments. However, 80% of the mutational data labelled with experimental outcomes from the same experiments it is tested on as well as fine-tuning with specific families as input. Previous neural language models^54-56^ are so far not suitable for mutation effect prediction for sequences without extensive experimental data or sequences with high variability, such as the complementarity-determining regions (CDRs) of antibody variable domains. By contrast, a fully unsupervised, alignment-free generative model of functional sequences is therefore desirable for the design of efficient nanobody libraries.

We then trained our validated statistical method on naïve nanobody repertoires^57^ as naïve antibody repertoires have been shown to have functional sequences with capacity to target diverse antigens^58^ and used it to generate probable sequences. In this manner we designed a sequence library that is 1000-fold smaller than state-of-art synthetic libraries but has an almost two-fold higher expression level, from which we identified a candidate binder for affinity maturation. A well designed library can also be used in continuously evolving systems^59^ to combine the hypermutation and affinity maturation processes of living organisms in a single experiment. Smart library design opens doors to more efficient search methods of nanobody sequence space for rapid discovery of stable and functional nanobodies.

## Results

### An autoregressive generative model of biological sequences

Protein sequences observed in organisms today result from mutation and selection for functional, folded proteins over time scales of a few days to a billion years. Generative models can be used to parameterize this view of evolution. Namely, they express the probability that a sequence ***x*** would be generated by evolution as *p*(***x***|***θ***), where parameters *θ* capture the constraints essential to functional sequences. An autoregressive model is one that makes a prediction in a time series (or sequence) using the previous observations. In our context, this means predicting the amino acid in a sequence using all of the amino acids that come before it. With the autoregressive model, the probability distribution *p*(***x***|***θ***) can be decomposed into the product of conditional probabilities on previous characters along a sequence of length L (**Supplementary Fig. 1**) via an autoregressive likelihood:

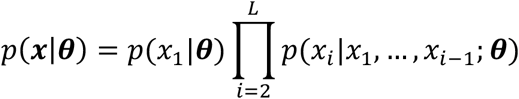

**Figure 1.**
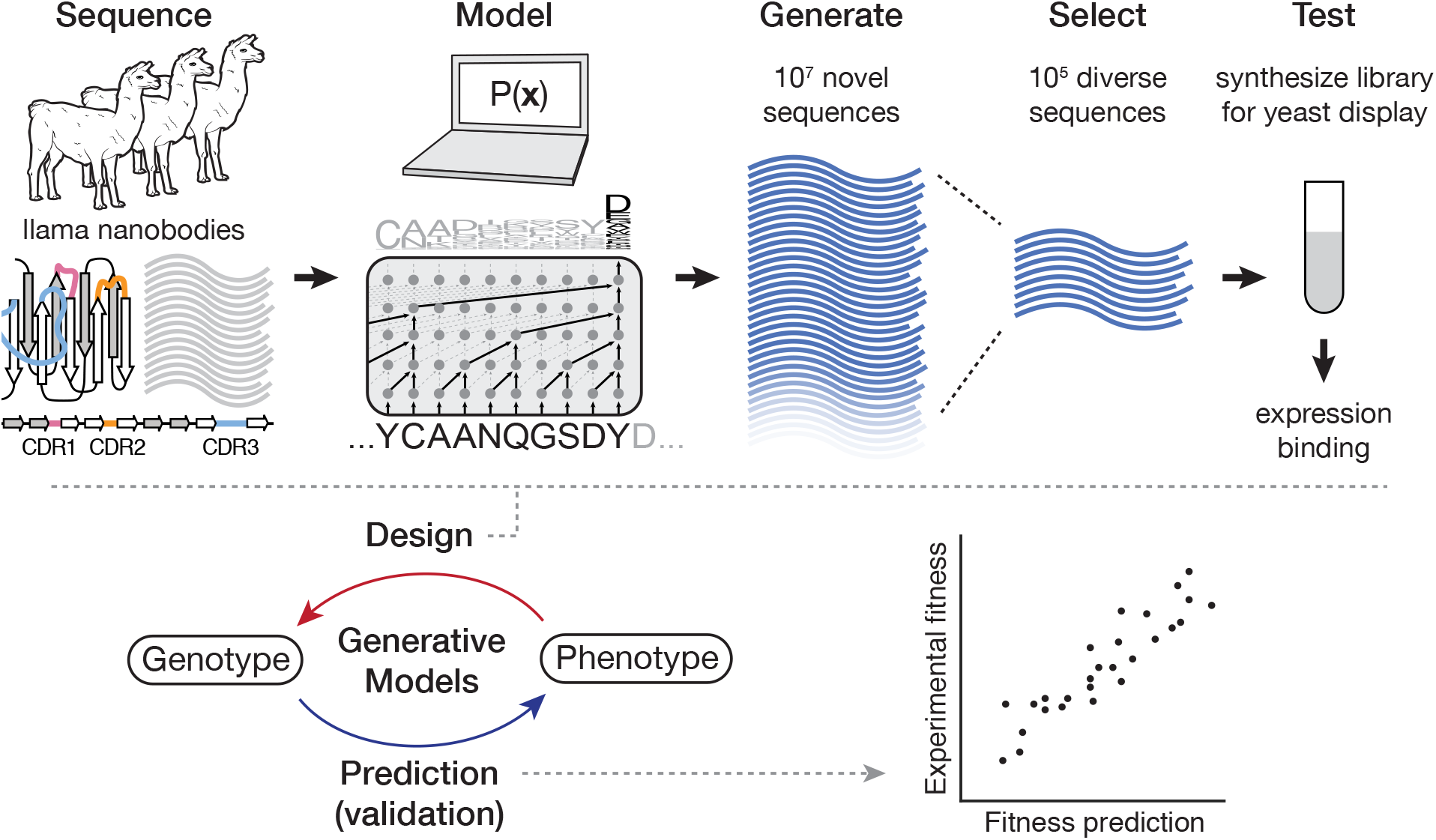
Autoregressive models of biological sequences can learn the genotype-phenotype map for both prediction and design. From natural sequences in a naïve llama repertoire^57^, the autoregressive model can learn functional constraints by predicting the likelihood of each residue in the sequence conditioned on preceding residues. We then use these constraints to generate millions of novel nanobody sequences—as many can be generated as desired. Of these designed sequences we select hundreds of thousands of diverse sequences, synthesize a library, and screen for expression and binding. We also validate the model on mutation effect prediction tasks of deep mutational scans including the effects of multiple insertions and deletions, and the thermostabilities of highly variable nanobody sequences.

Many different neural network architectures can model an autoregressive likelihood, including attention-based models^60^ and recurrent neural networks^61^. However, we encountered exploding gradients^62^ during training on long sequence families with LSTM^63^ or GRU^64^ architectures. Instead, we parameterize this process with dilated convolutional neural networks (**Supplementary Fig. 1**), which are feed-forward deep neural networks that aggregate long-range dependencies in sequences over an exponentially large receptive field^65-67^ (See Methods). The model is tasked with predicting an amino acid at some position in the sequence given all the previous amino acids in the sequence, i.e. forward language modeling. The causal structure of the model allows for efficient training to a set of sequences, inference of mutation effects, and sampling of new sequences. By learning these sequential constraints, the model can be directly applied to generating novel, fit proteins, one residue at a time. The autoregressive nature of this model obviates the need for a structural alignment and opens doors for application to modeling and design of previously challenging sequences such as non-coding regions, antibodies, and disordered proteins.

### The autoregressive model predicts experimental phenotype effects from sequences

In order to gain confidence in the new model for generating designed sequences, we first tested the ability of our new model to capture the dependencies between positions by testing the accuracy of mutation effect prediction. Somewhat surprisingly, unsupervised, generative models trained only on evolutionary sequences are proving the most accurate for predicting the effect of mutations when compared to large datasets of experimentally measured mutation effects^30, 34^, and they avoid the risk of overfitting that can occur as a result of circularity in supervised methods^68^. We compared the accuracy of this new, non-alignment-based model to state-of-art methods for a benchmark set of 40 deep mutational scans across 33 different proteins, totaling 690,257 individual sequences (**Supplementary Table 1**).

The autoregressive model was first fit to each family of protein sequences and then we used the log-ratio of likelihoods of individual sequences to predict mutation effects:

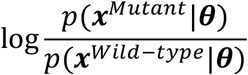

which estimates the plausibility of mutant sequence ***x***^*Mutant*^ relative to its wild-type, un-mutated counterpart, ***x***^*wild-type*^. This log-ratio has been shown to be predictive of mutation effects^29, 30^. Importantly, this approach is fully unsupervised: rather than learning from experimental mutation effects, we can learn evolutionary constraints using only the space of natural sequences. We benchmark the model predictions against the deep mutational scan experiments and compare the Spearman’s rank correlation to state-of-art models trained on alignments of the same sequences. The autoregressive model is able to consistently match or outperform a model with only site-independent terms (30/40 datasets) and the EVmutation model^29^ that includes dependencies between pairs of sites (30/40 datasets); it performs on par with the state-of-the-art results of DeepSequence^30^ (19/40 datasets, average difference in rank correlation is only 0.09); and it outperforms the supervised Envision model^31^ for 6/9 of the datasets tested (**Fig. 2a**; **Supplementary Figs. 2,3**). Previously published benchmarks^29^ demonstrate the higher accuracy of the probabilistic models, EVmutation compared to SIFT and PolyPhen, and recent work demonstrates that DeepSequence outperforms all currently available methods when measured against experimental mutation scans^34^. These benchmarks, taken together with our previous benchmarks^29^ and evidence from independent assessments^34^, show that our autoregressive model outperforms all methods including supervised and performs on par with our own state-of-art alignment-based method^30^ for single mutation effect prediction, providing us with the confidence to use the model for sequence design.

**Figure 2.**
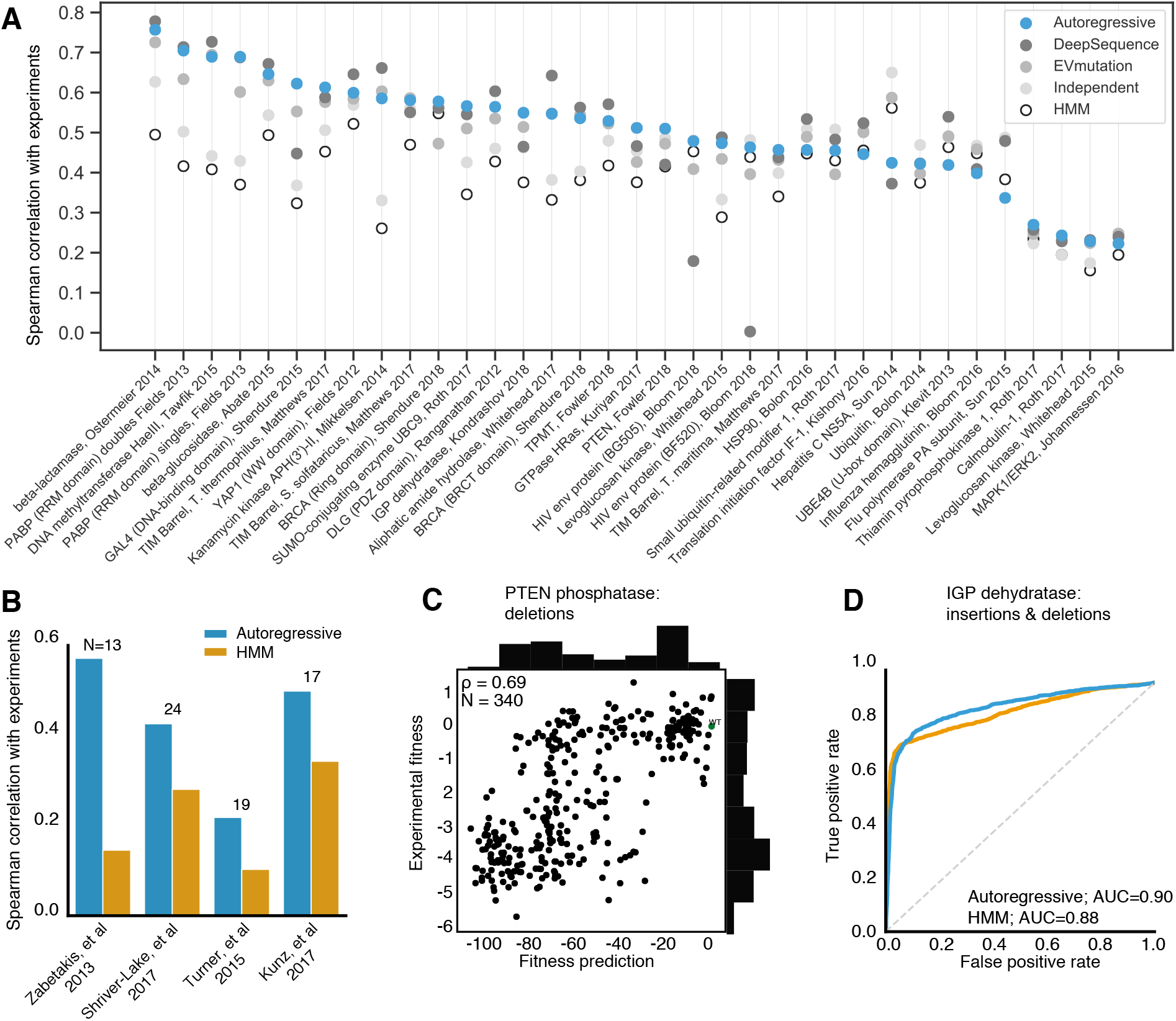
Validation of the autoregressive model in learning the genotype to phenotype map. The model accurately predicts fitness of biological sequences of various lengths. **a**. Even without using alignments, the autoregressive model can competitively match mutation effect prediction accuracies of state-of-art alignment-dependent models, such as conservation, evolutionary couplings, and DeepSequence. Additionally, the mutation effect prediction accuracies improves upon HMM model accuracies. Without using alignments, the autoregressive model matches alignment-dependent state-of-art missense mutation effect prediction (DeepSequence^30^) for 40 different deep mutational scan experiments. Three datasets show significant improvement with the autoregressive model: HIV env (BF520), HIV env (BG505), and GAL4 DNA-binding domain. **b**. The autoregressive model can learn from natural sequence repertoires of llama nanobodies to predict the thermostability of llama nanobody sequences with variation in the framework and complementarity determining regions with greater accuracy than hidden Markov models^74^. The number of llama nanobody sequences from each study is shown above each pair of bars. **c**. Fitness predictions for single deletions in PTEN phosphatase compared with measured experimental fitness is accurate, with a Spearman correlation of 0.69. **d**. Accurate prediction of binary fitness for IGP dehydratase with a range of insertions, deletions, and missense mutations.

**Figure 3.**
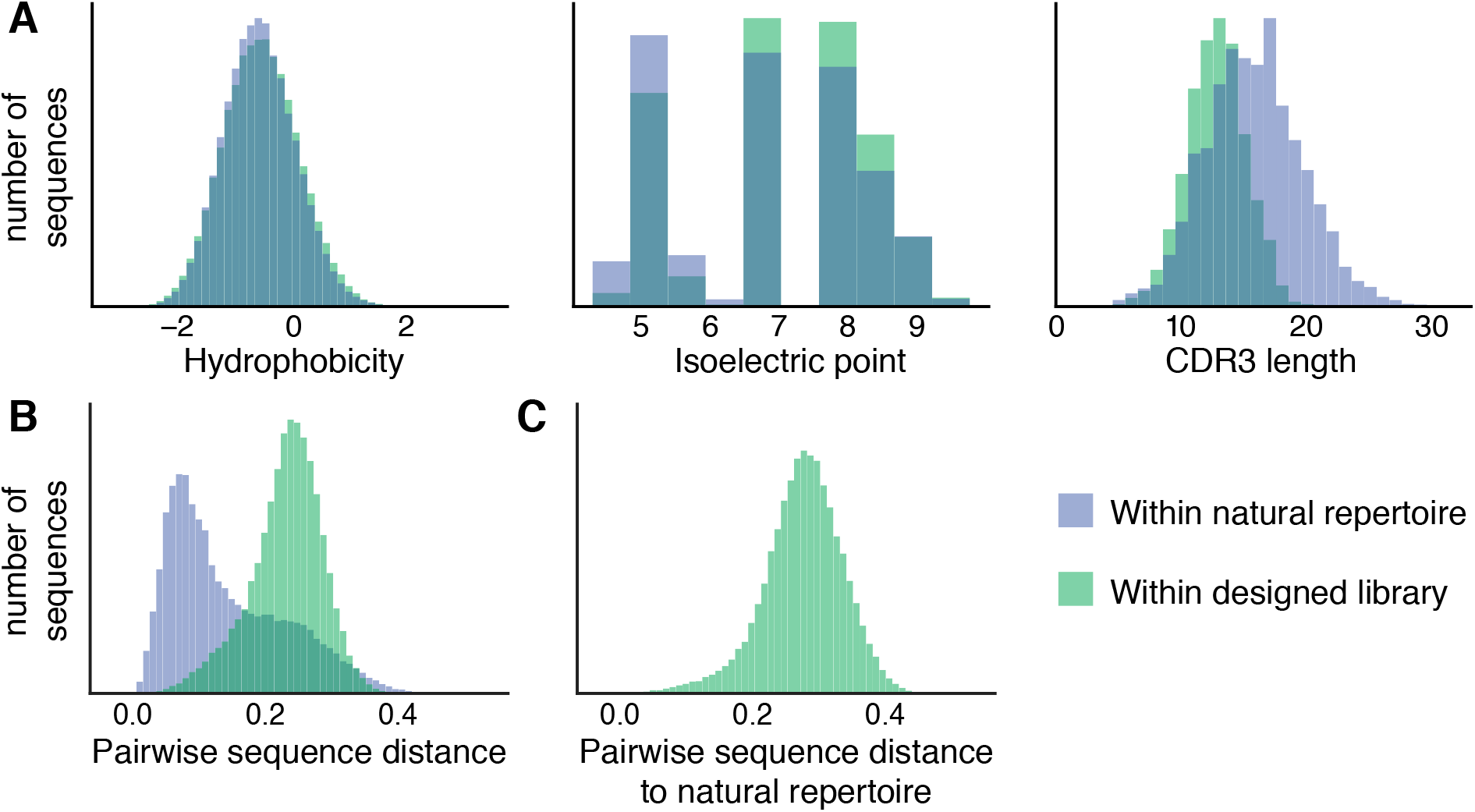
The designed library has comparable biochemical property distributions and improved diversity to the natural llama repertoire. **a**. Conditioned on the framework-CDR1-CDR2 sequence, a diverse set of CDR3 sequences are generated and selected. These CDR3 sequences are similar to the natural repertoire in their distributions of hydrophobicity^104^ and isoelectric point^105, 106^, while having shorter length distributions due to selection strategies in the final library construction. **b**. The designed library contains more diversity in sequences than the natural repertoire as evidenced by the larger cosine distance to its nearest neighbor. **c**. Each sequence in the designed library is diverse from any sequence seen in the natural repertoire, indicating that we have learned fit sequence constraints but are traversing previously unexplored regions of sequence space.

As with previous models that use evolutionary sequences, the accuracy of mutation effect prediction increases with increasing numbers of non-redundant sequences, as long as there is coverage of the length, tested here across eight of the protein families for four sequence depths (**Supplementary Fig. 4, Supplementary Table 2**). Interestingly, the accuracy of effect predictions against the aliphatic amidase mutation scan are remarkably robust even with a low number of training sequences—123 non-redundant sequences provide the same accuracy as 36,000—suggesting that there is more to learn about the relationship between evolutionary sampling and model learning. For now, we advise a conservative Meff/L (number of effective sequences normalized by length) requirement of 5 in order to sample enough diversity.

**Figure 4.**
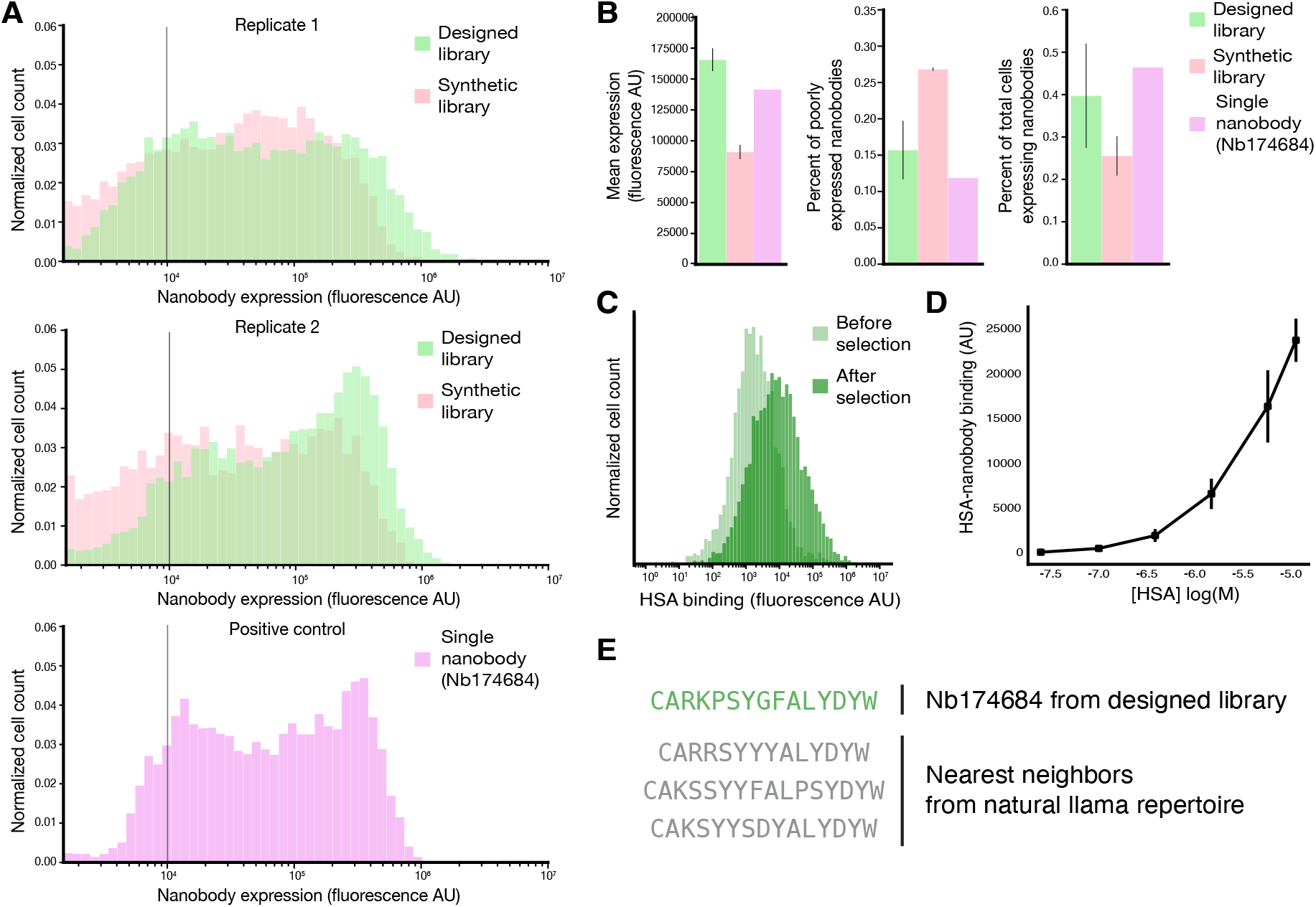
The designed library contains stable and functional nanobody sequences that are well expressed and can bind target antigens. **a**. Fluorescence distributions of cells expressing nanobodies comparing the synthetic combinatorial library and our designed library in two biological replicate experiments as well as a control experiment of a single, well-expressed nanobody clone (Nb174684). The distributions of the designed library are consistently right-shifted compared to the combinatorial library and resemble the control nanobody. **b**. Compared to the combinatorial library, the designed library has almost double the mean expression level (left panel, 166,193 AU compared to 92,183 AU), nearly half the fraction of poorly expressed nanobodies (of cells expressing nanobodies) (middle panel, 15.4% compared to 25.7% of clones with less than 10,000 AU indicated as a grey bar in panel **a**), and one and a half times the fraction of total cells that express nanobodies (right panel, 39.6% compared to 25.1%). The thresholds for determining the proportion of total cells expressing nanobodies were found by identifying the local minima on the distributions and are displayed in **Supplementary Fig. 10**. Values displayed on the bar graphs are means of the two replicates and the standard deviations are shown as error bars. There is only one replicate for the control experiment of the single nanobody clone. **c**. Fluorescence distributions of nanobodies bound to HSA shows a rightward shift after screening and selection, indicating a successful enrichment of binders to the target antigen. **d**. On-yeast binding assay of Nb.174684, an HSA binder identified from the designed library with moderate binding affinity. Error bars represent standard deviations in measurements at each concentration of HSA. **e**. CDR3 sequence of binder Nb.174684 and the sequences of the nearest neighbors from the natural llama repertoire that was used to train the autoregressive model.

Because the autoregressive model is not dependent on alignments, we can now learn mappings of sequences of high variability and diverse lengths for which meaningful alignments are difficult or non-sensical to construct, such as antibody and nanobody sequences. The autoregressive model was thus also validated on nanobody thermostability measurements to test whether we could learn the sequence constraints of fit nanobodies, including the highly variable regions. To do so, we fit the autoregressive model to a set of ∼1.2 million natural llama nanobody sequences^57^. Sequence likelihoods from this trained model are expected to reflect nanobody fitness, i.e., the multiple convolved aspects that nanobodies are selected for *in vivo*, including thermostability, expression, and potentially low polyreactivity. Using this model, we find that the log-probability fitness calculations predict the thermostability of unseen llama nanobody sequences from four different stability experiments^69-72^ (**Fig. 2b, Supplementary Fig. 5, Supplementary Table 3**). These experiments span a wide range of mutation types, lengths, and sequence diversity. The autoregressive model consistently outperforms a hidden Markov model (HMM, hmmer3)^73, 74^ in predicting the relationship between sequence and thermostability of nanobodies.

Previous alignment-dependent generative models are constrained to predicting the effects of missense mutations. However, in-frame insertions and deletions can also have large phenotypic consequences for protein function, yet these changes have proved difficult to model. We compare the fitness predictions calculated as log probabilities by the autoregressive model to experimental assays for the fitness of mutated biomolecules, using rank correlation (ρ) for quantitative measurements and area under the receiver-operator curve (AUC) for binary fitness categorization, identifying the two groups with a two-component Gaussian mixture model. The model is able to capture the effects of single amino acid deletions on PTEN phosphatase^75^ (ρ=0.69, N=340, HMM ρ=0.75; PROVEAN ρ=0.7; **Fig. 2c**) and multiple amino acid insertions and deletions in imidazoleglycerol-phosphate (IGP) dehydratase^76^ (AUC=0.90, N=6102, HMM AUC=0.88; **Fig. 2d, Supplementary Table 4**). Here we use the AUROC metric for IGP dehydratase as the experimental data are bimodal with a large fraction at zero fitness. While PROVEAN^77^ predicted the effect of single PTEN deletions comparably to our model, it fails to predict the effect of multiple insertions, deletions, and substitutions as were tested in IGP dehydratase and it cannot generate new sequences. Three additional insertion and deletion mutation scan fitness predictions are included in the supplement: yeast snoRNA (ρ=0.49), beta lactamase (ρ=0.45), and p53 (ρ=0.035; **Supplementary Fig. 6**). Predicting the effects of indels also has clinical significance: the four different single amino acid deletions annotated as pathogenic by Clinvar^78^ in two cancer genes, BRCA1 and P53, and one Alzheimer’s-linked gene, APOE, are in the bottom 25^th^ percentile of predicted deletion effect distributions (**Supplementary Fig. 7**). Other indels that are predicted to be highly deleterious by the autoregressive model may be of clinical interest for experimental study of pathogenicity. We expect that the autoregressive model can predict mutation effects in disordered and low-complexity sequences. As a proof-of-concept, we have provided an *in silico* mutation scan of the human tau protein, which contains regions of low complexity and is strongly associated with neurodegenerative diseases, (**Supplementary Fig. 8**). Our mutation effect prediction distinguishes between 40 pathogenic and 10 non-pathogenic mutations (two-tailed independent t=-4.1, P=0.001, AUC=0.86) that were collected from the Alzforum database^79^.

### Generating an efficient library of functional nanobodies

Screening large, high-throughput libraries of antibodies and nanobodies in vitro has become increasingly prevalent because it can allow for rapid identification of diverse monoclonal binders to target antigens. However, these synthetic libraries contain a large fraction of non-functional nanobody sequences. Natural nanobody sequences are selected against unfavorable biochemical properties such as instability, poly-reactivity, and aggregation during affinity maturation^6^. Similarly to nanobody thermostability prediction, we sought to learn the constraints that characterize functional nanobodies by fitting the autoregressive model to a set of ∼1.2 million nanobody sequences from the immune repertoires of seven different naïve llamas^57^. Using this trained model and conditioning on the germline framework-CDR1-CDR2 nanobody sequence, we then generate over 10^7^ fit sequences, generating one amino acid at a time based on the learned sequential constraints. As nanobody CDR3s often contact the framework in 3D, conditioning in this way allows the model to learn any resulting constraints on the CDR3 sequence and incorporate them during generation. We remove sequences that do not end with the final beta strand of our nanobody template, duplicate sequences, and CDR3s likely to suffer post-translational modification to obtain ∼3.7 million sequences (**Supplementary Table 5**). From these, we select 185,836 highly diverse CDR3 sequences for inclusion in our designed library. We compare our designed library to a state-of-art synthetic library^8^, which was constructed combinatorically based on position specific amino acid frequencies of nanobody sequences with crystal structures in the PDB database. This library contains CDR3 sequences that have a similar distribution of biochemical properties as the naïve llama immune repertoire (Methods; **Fig. 3a**). The distribution of hydrophobicity and isoelectric points are similar to the natural llama repertoire even though explicit constraints on these properties were never imposed during generation or selection of sequences for the designed library. The lengths of the CDR3 sequences in the designed library are shorter than the natural repertoire; this is due to the strategy of choosing cluster centroids during selection of the 10^5^ sequences and can be adjusted by changing the sampling method. Longer CDR3s may also be attained by allowing interloop disulfide bridges that stabilize longer CDR3s in some VHH domains^80^; this would require a different nanobody template and ideally camel or dromedary nanobody repertoires. The sequences in the designed library are extremely diverse and are more distant from each other than sequences in the natural repertoire (**Fig. 3b**), while maintaining nearly as much diversity as an equivalent sample of a combinatorial synthetic library^8^ (**Supplementary Fig. 9**). Additionally, we are exploring new regions of sequence space because the generated sequences in the designed library are diverse from the naïve repertoire (**Fig. 3c**).

Using these designed CDR3 sequences, a nanobody library was constructed using our yeast-display technology for experimental characterization alongside a combinatorial synthetic nanobody library^8^. The designed library had more length diversity and a longer CDR3 median length (13) than the synthetic library (12) (**Supplementary Fig. 9**), while the synthetic library included designed diversity in specific residues of the CDR1 and CDR2. Individual nanobody sequences were expressed on the surface of yeast cells, allowing for rapid sorting of nanobody clones based on expression and/or binding levels. Upon induction, the designed nanobody library contained 1.5 times higher proportion of cells expressing and displaying nanobodies on their cell surface than the synthetic nanobody library (**Fig. 4a**,**b, Supplementary Fig. 10**). In the designed library, we can also see a clearer separation of cells expressing nanobodies and those that are not.

Of cells expressing nanobodies, the mean nanobody display levels from the designed library is almost twice the level of the previous library (**Fig. 4a**,**b**). Furthermore, the designed library had nearly half the fraction of poorly expressed nanobodies (cells with fluorescence below 10,000 AU) as compared to the synthetic library (**Fig. 4a**,**b**) as well as a significant increase in the fraction of highly expressed nanobodies as can be seen in the upper limits in the respective expression distributions (**Fig. 4a, Supplementary Fig. 10**). Expression experiments were performed with two replicates in addition to a single control experiment of yeast expressing a single well-behaved nanobody clone (Nb. 174684). These experimental results demonstrate that with the autoregressive model trained on natural llama nanobody sequences, we successfully designed a smart library consisting of a higher proportion of stable, well-expressed nanobodies. With this small designed library, we selected nanobody sequences that bound to human serum albumin (HSA) using fluorescence activated cell sorting (FACS) (**Fig. 4c**), from which we were even able to identify weak to moderate binders—the strongest binder has a predicted *K_d_* of 9.8 *μM* (**Fig. 4d**). This experiment is a proof-of-concept that this small library contains antigen-binding sequences that can be starting points for affinity maturation to identify strong binders. Though not explicitly designed to minimize poly-reactive nanobody sequences, training on a naïve llama repertoire, which presumably contain a moderate proportion of poly-reactive sequences^81-87^, the designed library shows similar levels of poly-reactivity to the synthetic library, which had been designed according to a small set of highly specific nanobodies (**Supplementary Fig. 11**). These results indicate that we have successfully designed an efficient library containing a high proportion of promising diverse, stable, specific, and sensitive nanobody sequences.

## Discussion

Here we show how neural network-powered generative autoregressive models can be used to model sequence constraints independent of alignments and design novel functional sequences for previously out of reach applications such as nanobodies. The capability of these models is based on demonstrated state-of-the-art performance and on an extended range of applicability in the space of sequences. In the particular version in this paper, we validated our model first on deep mutational scan data, with on par performance with the best currently available model^29-31, 34, 77^, and demonstrated application to examples for which robust alignments cannot be constructed, such as sequences with multiple insertions, deletions, and substitutions, and cases for which protein structures and experimental data are not available. As for comparison with a potentially competing alignment-free model, while we do not discount the utility of semi-supervised methods (exploiting mutation effect-labeled experimental data), great care must be taken in the way the split between training and test is conducted to evaluate the true generalizability of the method. For instance, randomized subsets excluded from training will still be learned from the labeled data in a way that is not generalizable to required predictions for other proteins^53,88,89^. Our model is not subject to these limitations as its training is fully unsupervised.

Due to their flexibility, deep autoregressive models could also open the door to new opportunities in biological sequence analysis and design. Unlike alignment-based techniques, since no homology between sequences is explicitly required, generative models with autoregressive likelihoods can be applied to variants with insertions and deletions, disordered proteins, multiple protein families, promoters and enhancers, or even entire genomes. Specifically, prediction of insertions and deletions and mutation effects in disordered regions has been a difficult research area, despite their prevalence in human genomes. Disordered regions are enriched in disease-associated proteins, so understanding variant effects will be important in understanding the biology and mechanism of genes indicated in cardiovascular, cancer, and neurodegenerative diseases. For example, classical tumor suppressor genes, such as p53, BRCA1, and VHL, and proteins indicated in Alzheimer’s disease, such as Tau, have long disordered regions where these models may prove particularly useful.

With this model, we designed a smart, diverse, and efficient library of fit nanobody sequences for experimental screening against target antigens. Designing individual hypervariable CDR sequences that make up a library of diverse, functional, and developable nanobodies allows for much faster and cheaper discovery of new therapeutics, minimizing both library waste and necessary experimental steps. Our streamlined library (1000-fold smaller than combinatorial synthetic libraries) enables rapid, efficient discovery of candidate nanobodies, quickly providing a starting point for affinity maturation to enhance binding affinity. In combination with a continuous evolution system, candidate binders from the designed library have been identified and affinity matured after only a few rounds of selection with a single experiment^90^. As the cost to synthesize sequences decreases, the demand for methods that can design highly optimized and diverse sequences will increase as compared to constructing libraries via random or semi-random generation strategies.

A challenge of using synthetic libraries is the poly-reactivity of many sequences that *in vivo*, would be cleared by an organism’s immune system. Naïve llama repertoires also contain poly-specific sequences, so training a model on sequences from mature or memory B cell repertoires may provide information on how to improve library design in the future and minimize the poly-reactivity of the designed library sequences. Multi-chain proteins such as antibodies present an additional challenge that multiple domains must be designed together. Models incorporating direct long-range interactions such as dilated convolutions or attention may identify the relevant dependencies between domains, even when the domains simply concatenated and generated sequentially. Paired antibody chains are more challenging to sequence than nanobodies, but more repertoires are becoming available^91^. Beyond antibody and antibody fragment libraries, this method is translatable to library design for any biomolecule of interest, including disordered proteins.

Our model is the first alignment-free method demonstrating state-of-art mutation effect prediction without experimental data and applied to at scale to design of protein sequences. New developments in machine learning will enhance the power of such autoregressive models and incorporating protein structural information may further improve the capacity to capture long-range dependencies^92^ for these applications. The addition of latent variables could also allow for targeted design of high affinity and specificity sequences to a desired target antigen^56, 93-95^. Conversely, we also anticipate better exploration of broader spans of sequence space for generation, either by exploiting variance explained by latent variables^96^ or diverse beam search strategies^97^. With the increased number of available sequences and growth in both computing power and new machine learning algorithms, autoregressive sequence models may enable exploration into previously inaccessible pockets of sequence space.

## Methods

### Model

Sequences are represented by a 21-letter alphabet for proteins or 5-letter alphabet for RNAs, one for each residue type and a ‘start/stop’ character. Training sequences are weighted inversely to the number of neighbors for each sequence at a minimum identity of 80%, except for viral families, where a 99% identity threshold was used, as was done previously^30^. Sequence sets are derived from alignments by extracting full sequences for each aligned region; sequence identities, boundaries, and weights are the only information provided to the model by alignments. The log-likelihood for a sequence is the sum of the cross-entropy between the true residue at each position and the predicted distribution over possible residues, conditioned on the previous characters. Since we encountered exploding gradients^62^ during training on long sequence families with LSTM^63^ or GRU^64^ architectures, we parameterize an autoregressive likelihood with dilated convolutional neural networks (**Supplementary Fig. 1**). These feed-forward deep neural networks aggregate long-range dependencies in sequences over an exponentially large receptive field^65-67^. Specifically, we use a residual causal dilated convolutional neural network architecture with 6 blocks of 9 dilated convolutional layers and both weight normalization^98^ and layer normalization^99^, where the number of blocks and layers were chosen to cover protein sequences of any length. To help prevent overfitting, we use L2 regularization on the weights and place Dropout layers (p = 0.5) immediately after each of the 6 residual blocks^100^. We use a batch size of 30 for all sequence families tested. Channel sizes of 24 and 48 were tested for all protein families, and channel size 48 was chosen for further use. Six models are built for each family: three replicates in both the N-to-C and C-to-N directions, respectively. Each model is trained for 250,000 updates using Adam with default parameters^101^ at which point the loss had visibly converged, and the gradient norm is clipped^62^ to 100.

### Data collection

40 datasets which include experimental mutation effects, the sequence families, and effect predictions were taken from our previous publication^30^ and 5 datasets that include indels and nanobody thermostability data were added for this work (references and data in **Supplementary Table 4** and **Extended Data**). For new mutation effect predictions such as the indel mutation scans, sequence families were collected from the UniProt database in the same procedure as described in previous published work^30^. Pathogenic muations for the Tau protein were downloaded from the Alzforum database^79^. The naïve llama immune repertoire was acquired from^57^. Due to the large number of sequences in the llama immune repertoire, sequence weights were approximated using Linclust^102^ by clustering sequences at both 80% and 90% sequence identity thresholds.

### Nanobody library generation

Using the N-to-C terminus model trained on llama nanobody sequences, we generated 33,047,639 CDR3 sequences by ancestral sampling^61^, conditioned on the germline framework-CDR1-CDR2 sequence and continued until generation of the stop character. Duplicates of the training set or generated sequences and those not matching the final beta strand of our nanobody template were excluded. CDR3 sequences were also removed if they contained glycosylation (NxS and NxT) sites, asparagine deamination (NG) motifs, or sulfur-containing amino acids (cysteine and methionine), resulting in 3,690,554 sequences.

From this large number of sequences, we then sought to choose roughly 200,000 CDR3 sequences that are both deemed fit by the model and as diverse from one another as possible to cover the largest amount of sequence space. First, we featurized these sequences into fixed length, L2 normalized k-mer vectors with k-mers of size 1, 2, and 3. We then used BIRCH clustering^103^ to find diverse members of the dataset in O(n) time. We used a diameter threshold of 0.575, resulting in 382,675 clusters. K-mer size and BIRCH diameter threshold were chosen to maximize the number of clusters within a memory constraint of 70 GB. From the cluster centroids, we chose the 185,836 most probable sequences for final library construction.

### Construction of nanobody library

FragmentGENE_NbCM coding for the nanobody template was amplified with oligonucleotides NbCM_pydsF2.0 and NbCM_pydsR and then cloned into the pYDS649 yeast-display plasmid^8^ using HiFi Mastermix (New England Biolabs). The original NotI site in pYDS649 was then removed by amplification with primers NotI_removal_1F and Pyds_NbCM_cloning_R followed by cloning again into pYDS649 to generate the pYDS_NbCM display plasmid for the nanobody template.

An oligonucleotide library was synthesized (Agilent) with the following design ACTCTGT [CDR3] ATCGT where CDR3 is a sequence for one of the computationally designed clones. Two-hundred picomoles of the library was PCR amplified over 15 cycles with oligonucleotides Oligo_library_F and Oligo_library_R using Q5 polymerase (New England Biolabs). Amplified DNA was PCR purified (Qiagen) and ethanol precipitated in preparation for yeast transformation. 4.8 × 10^8^ BJ5465 (MATα ura352 trp1 leu2Δ1 his3Δ200 pep4::HIS3 prb1Δ1.6 R can1 GAL) yeast cells, grown to OD600 1.6, were transformed, using an ECM 830 Electroporator (BTX-Harvard Apparatus), with 2.4 µg of NotI digested pYDS_NbCM vector and 9.9 µg of CDR3 library PCR product yielding 2.7 × 10^6^ transformants. Library aliquots of 2.4 × 10^8^ cells per vial were frozen in tryptophan dropout media containing 10% DMSO.

### Characterization of nanobody library

Yeast displaying the computationally designed or combinatorial synthetic nanobody library^8^ were grown in tryptophan dropout media with glucose as the sugar source for one day at 30 °C and then passaged into media with galactose as the sole sugar source to induce expression of nanobodies at 25 °C. After two days of induction, one million cells from each library were stained with a 1:25 dilution of anti-HA AlexaFluor647 conjugated antibody (Cell Signaling Technology) in Buffer A (20 mM HEPES pH 7.5, 150 mM NaCl, 0.1% BSA, 0.2% maltose) for 30 minutes at 4 °C. After staining, cells were centrifuged, the supernatant was removed, and cells were resuspended in Buffer A for flow analysis with an Accuri C6 (BD Biosciences, **Supplementary Fig. 12**).

To find nanobody binders to human serum albumin (HSA) one round of magnetic-activated cell sorting (MACS) followed by two rounds of fluorescence-activated cell sorting (FACS) were performed on our yeast-displayed library of nanobodies. For MACS, 4 × 10^7^ induced cells were resuspended in binding buffer (20 mM HEPES pH 7.5, 150 mM NaCl, 0.1% ovalbumin) along with anti-fluorescein isothiocyanate (FITC) microbeads (Miltenyi) and FITC-labeled streptavidin for 35 min at 4°C and then passed through an LD column (Miltenyi) to remove binders to microbeads and streptavidin. Remaining yeast were centrifuged and resuspended in binding buffer and incubated with 500 nM streptavidin-FITC and 2 µM of biotinylated HSA for one hour at 4°C. Yeast were then centrifuged and resuspended in binding buffer containing anti-FITC microbeads for 15 min at 4°C before passing them into an LS column and eluting and collecting the bound yeast. For the first round of FACS, induced yeast were first stained with 1 µM of biotinylated HSA for 45 min at 4°C and then briefly stained with 500 nM of streptavidin tetramer along with antiHA-488 to assess expression levels. Both yeast stainings were performed in FACS buffer (20 mM HEPES pH 7.5, 150 mM NaCl, 0.1% ovalbumin, 0.2% maltose). 5 × 10^6^ yeast were sorted and 28,000 were collected and expanded for a second round of FACS. The second round of FACS was performed under the same conditions as the first and from 3.8 × 10^6^ sorted yeast 21,455 were collected. Nanobody Nb174684 was isolated from a screen of 36 clones for binding to HSA using a flow cytometer and then sequenced. In order to characterize binding of Nb174684, yeast displaying Nb174684 were stained with varying amounts of AlexaFluor 488 labeled HSA and fluorescence was analyzed with a flow cytometer.

### Oligonucleotides

FragmentGENE_NbCM:

GCTGCCCAGCCGGCGATGGCCCAGGTCCAACTTCAAGAATCAGGCGGGGGCCTGGTACAGGCAGGCGGTTCTCTTCGGCTGTCGTGTGCGGCAAGCGGATTTACATTCAGTAGCTACGCTATGGGCTGGTACCGTCAGGCACCGGGGAAAGAACGGGAATTTGTTGCTGCAATCTCTTGGAGCGGTGGGAGCACATATTATGCAGATTCCGTTAAAGGCAGATTCACGATCAGTCGCGATAACGCAAAAAATACAGTGTACTTACAAATGAACTCTTTGAAACCCGAAGACACCGCAGTCTATTACTGCGCGGCCGCTACTGGGGACAAGGCACCCAGGTGACTGTATCATCCCACCACCACCACCACCACTGA

NbCM_pydsF2.0:

GGTGTTCAATTGGACAAGAGAGAAGCTGACGCAGAAGTCCAACTTGTCGAATCAGGCGGGGGCCTGGTACAG

NbCM_pydsR: CGTAATCTGGAACATCGTATGGGTAGGATCCGGATGATACAGTCACCTGGGT

NotI_removal_1F: CAACCCTCACTAAAGGGCGTTCGCCATGAGATTCCCATCTATCTTCA

Pyds_NbCM_cloning_ R: CACCTGGGTGCCTTGTCCCCAGTA

## Acknowledgments

We would like to thank John Ingraham, members of the Marks and Sander labs, and Harvard Research Computing for their insight and feedback to our research.

## Author contributions

D.S.M., A.C.K., and A.J.R. conceived the project; A.J.R. constructed the model; A.J.R, J.-E.S., and A.W.K. designed and evaluated computational experiments for validation, prediction, and generation of sequences; A.M. compiled natural nanobody sequence data; C.M. constructed the library and performed experiments; J.-E.S., A.W.K., and C.M. analyzed the library experimental data; A.J.R., J.-E.S., A.W.K., C.M., A.C.K., and D.S.M. wrote the manuscript.

## Data and code availability

All data generated and analyzed during the study are available in this published article, its supplementary information files and on the github repository (https://github.com/debbiemarkslab/SeqDesign). All code used for model training, mutation effect prediction, sequence generation, and library generation is also available on the github repository.

## Supplementary Information

Supplementary Figures 1-12

Supplementary Tables 1-5

Extended Data 1-6

