## Supplementary Information for "Protein Design and Variant Prediction Using Autoregressive Generative Models"

**This file includes:**

Supplementary Figures 1 to 12
Supplementary Tables 1 to 5
Supplementary Note
Captions for Extended Data 1 to 6

### Supplementary Figures

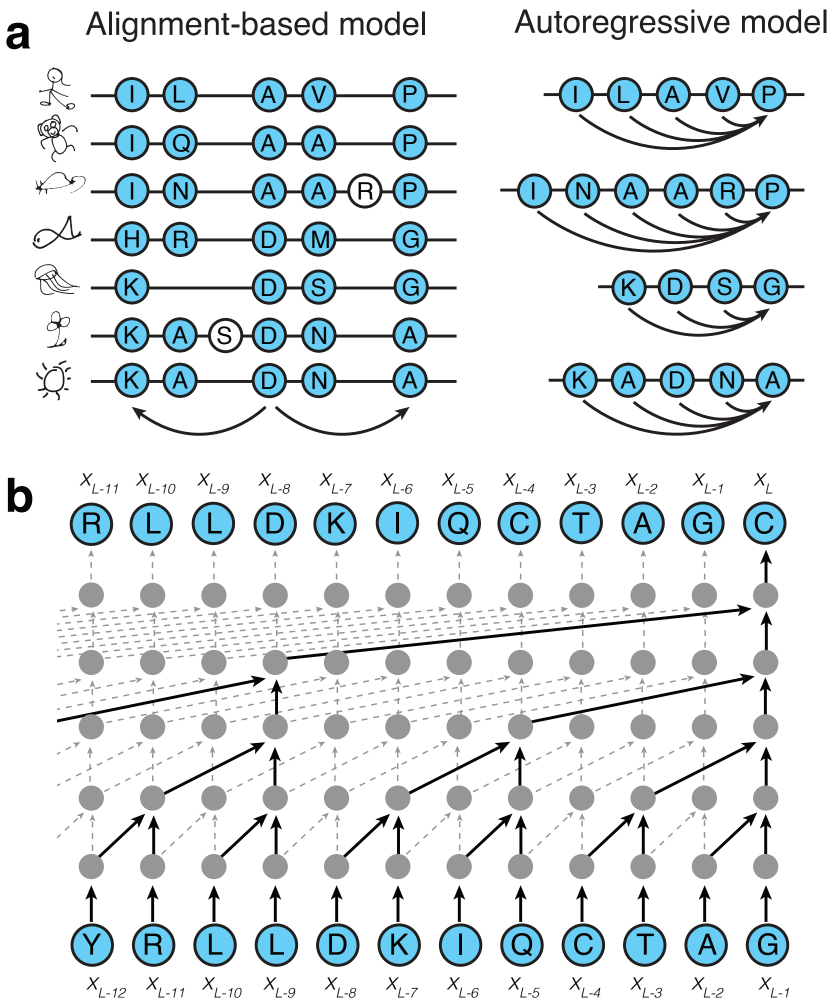


**Supplementary Fig. 1.** Autoregressive models of biological sequences. **a**. Instead of finding correlations between columns in a multiple sequence alignment (left), the autoregressive model predicts a residue given all the preceding positions (right). **b**. Causal dilated convolutions are used to model the autoregressive likelihood.

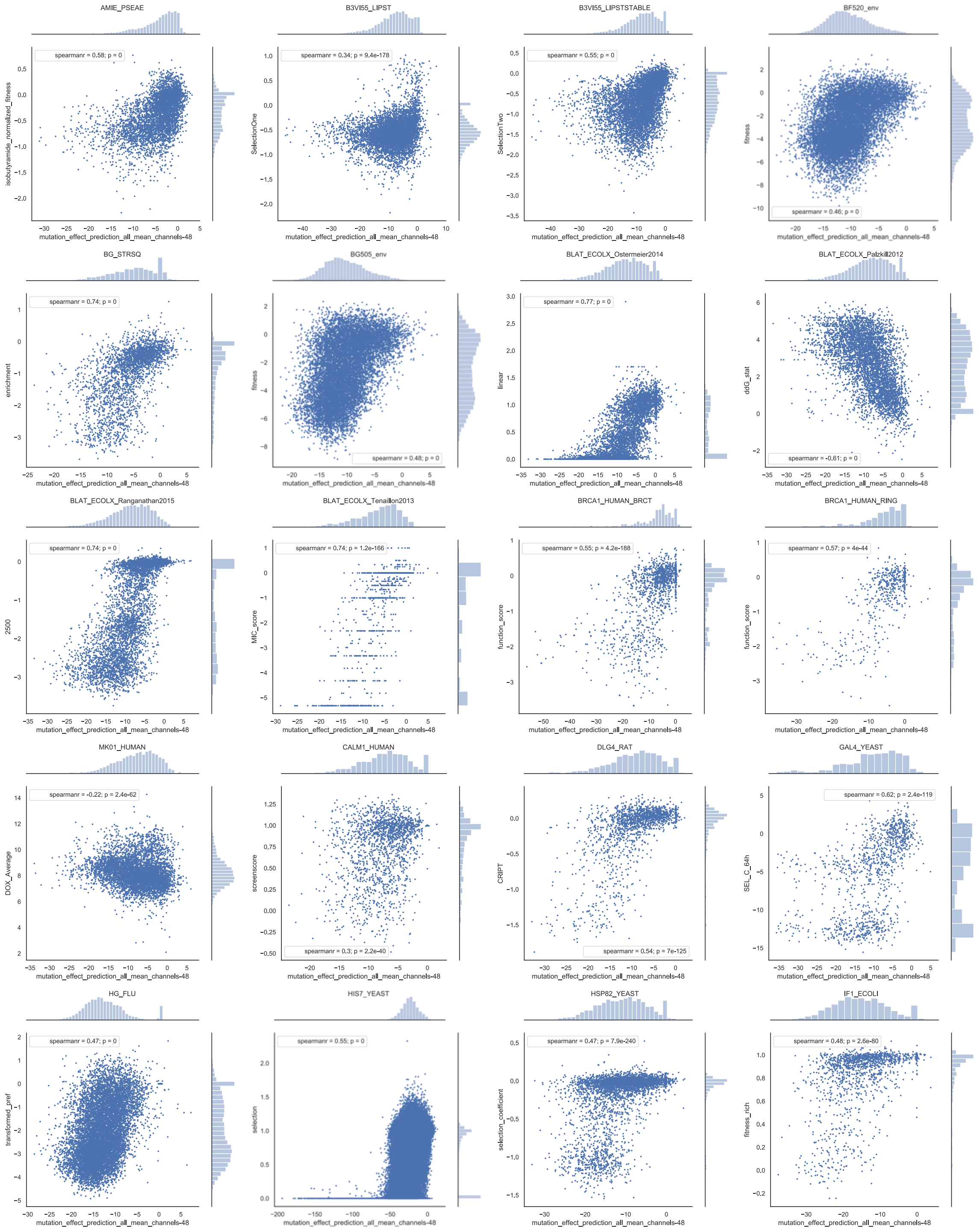
AMIE_PSEAE
spearmanr = 0.58; p = 0
isobutyramide_normalized_fitness
mutation_effect_prediction_all_mean_channels-48
B3VI55_LIPST
spearmanr = 0.34; p = 9.4e-178
SelectionOne
mutation_effect_prediction_all_mean_channels-48
B3VI55_LIPSTSTABLE
spearmanr = 0.55; p = 0
SelectionTwo
mutation_effect_prediction_all_mean_channels-48
BF520_env
spearmanr = 0.46; p = 0
fitness
mutation_effect_prediction_all_mean_channels-48
BG_STRSQ
spearmanr = 0.74; p = 0
enrichment
mutation_effect_prediction_all_mean_channels-48
BG505_env
spearmanr = 0.48; p = 0
fitness
mutation_effect_prediction_all_mean_channels-48
BLAT_ECOLX_Ostermeier2014
spearmanr = 0.77; p = 0
linear
mutation_effect_prediction_all_mean_channels-48
BLAT_ECOLX_Palzkill2012
spearmanr = -0.61; p = 0
ddG_stat
mutation_effect_prediction_all_mean_channels-48
BLAT_ECOLX_Ranganathan2015
spearmanr = 0.74; p = 0
2500
mutation_effect_prediction_all_mean_channels-48
BLAT_ECOLX_Tenaillon2013
spearmanr = 0.74; p = 1.2e-166
MIC_score
mutation_effect_prediction_all_mean_channels-48
BRCA1_HUMAN_BRCT
spearmanr = 0.55; p = 4.2e-188
function_score
mutation_effect_prediction_all_mean_channels-48
BRCA1_HUMAN_RING
spearmanr = 0.57; p = 4e-44
function_score
mutation_effect_prediction_all_mean_channels-48
MK01_HUMAN
spearmanr = -0.22; p = 2.4e-62
DOX_Average
mutation_effect_prediction_all_mean_channels-48
CALM1_HUMAN
spearmanr = 0.3; p = 2.2e-40
screenscore
mutation_effect_prediction_all_mean_channels-48
DLG4_RAT
spearmanr = 0.54; p = 7e-125
CRIPT
mutation_effect_prediction_all_mean_channels-48
GAL4_YEAST
spearmanr = 0.62; p = 2.4e-119
SEL_C_64h
mutation_effect_prediction_all_mean_channels-48
HG_FLU
spearmanr = 0.47; p = 0
transformed_pref
mutation_effect_prediction_all_mean_channels-48
HIS7_YEAST
spearmanr = 0.55; p = 0
selection
mutation_effect_prediction_all_mean_channels-48
HSP82_YEAST
spearmanr = 0.47; p = 7.9e-240
selection_coefficient
mutation_effect_prediction_all_mean_channels-48
IF1_ECOLI
spearmanr = 0.48; p = 2.6e-80
fitness_rich
mutation_effect_prediction_all_mean_channels-48

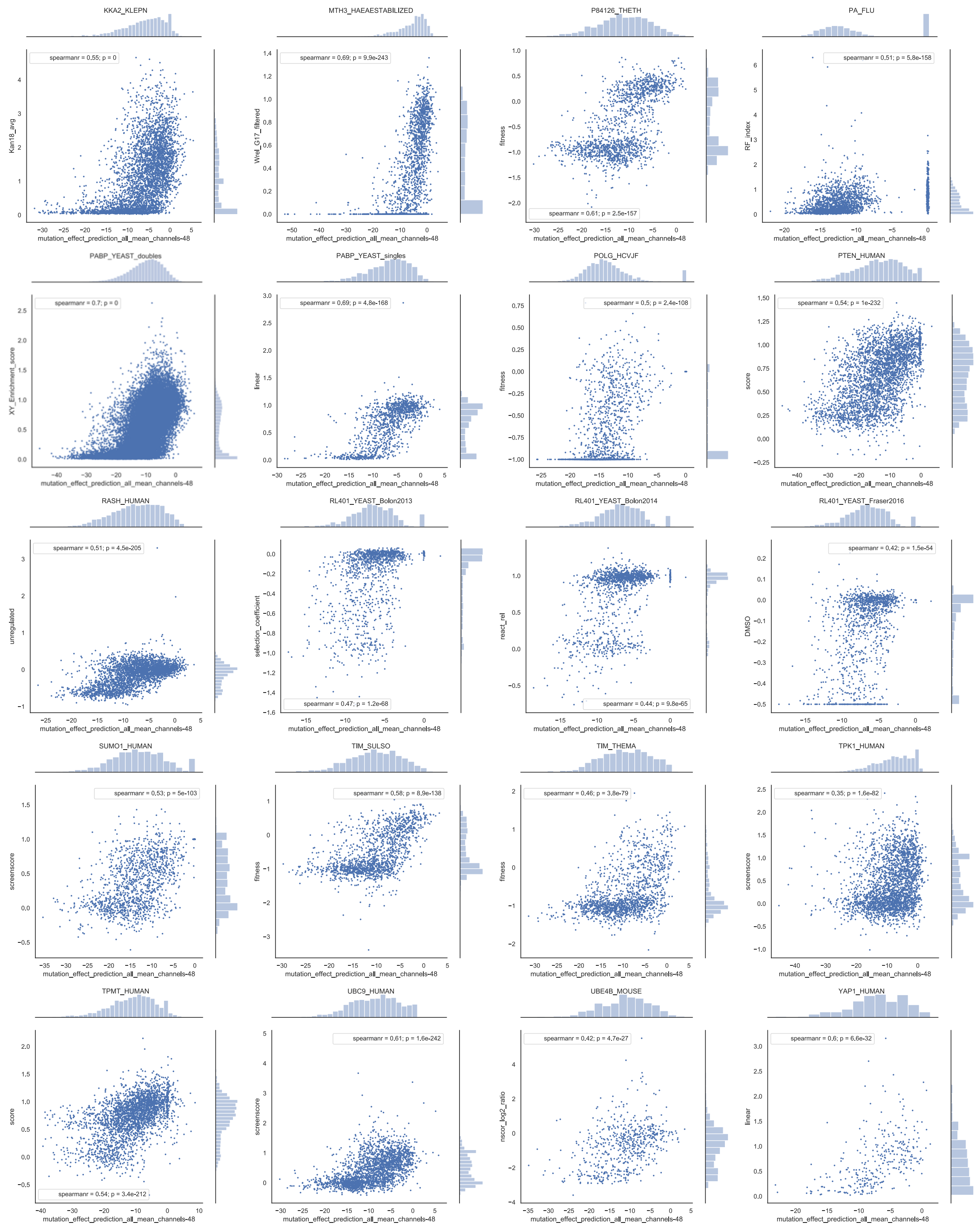


**Supplementary Fig. 2.** Individual scatterplots of the experimental results and the mutation effect prediction using the autoregressive model trained on each individual family of naturally occurring proteins. Experimental measures of fitness are on the y-axis and fitness predictions are on the x-axis.

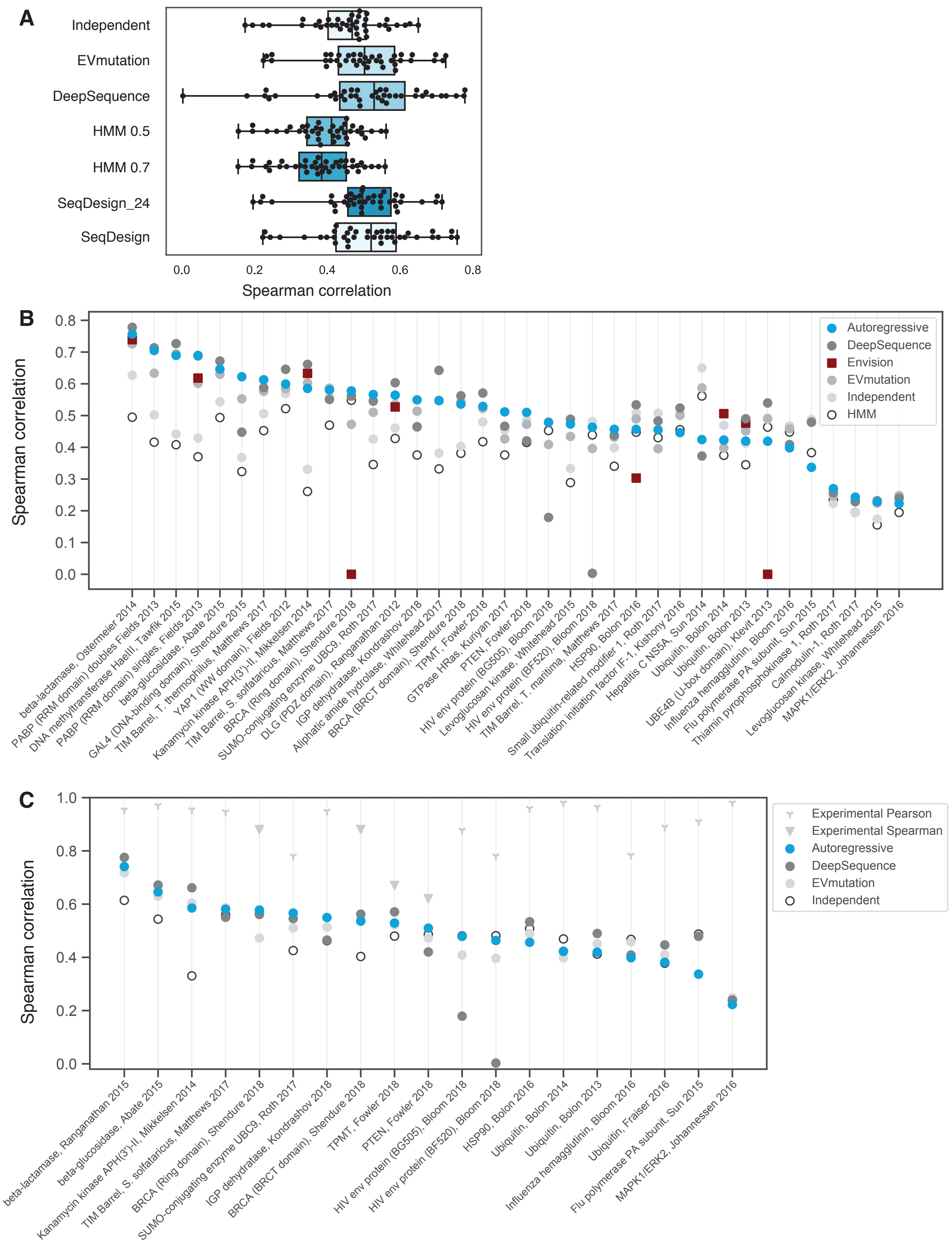
A
Independent
EVmutation
DeepSequence
HMM 0.5
HMM 0.7
SeqDesign_24
SeqDesign
0.0
0.2
0.4
0.6
0.8
Spearman correlation
B
Spearman correlation
0.8
0.7
0.6
0.5
0.4
0.3
0.2
0.1
0.0
Autoregressive
DeepSequence
Envision
EVmutation
Independent
HMM
beta-lactamase, Ostermeier 2014
PABP (RRM domain) doubles Fields 2013
DNA methyltransferase HaeIII, Tawfik 2015
PABP (RRM domain) singles, Fields 2013
beta-glucosidase, Abate 2015
GAL4 (DNA-binding domain), Shendure 2015
TIM Barrel, T. thermophilus, Matthews 2017
YAP1 (WW domain), Fields 2012
Kanamycin kinase APH(3')-II, Mikkelsen 2014
TIM Barrel, S. solfataricus, Matthews 2017
BRCA (Ring domain), Shendure 2018
SUMO-conjugating enzyme UBC9, Roth 2017
DLG (PDZ domain), Ranganathan 2012
IGP dehydratase, Kondrashov 2018
Aliphatic amide hydrolase, Whitehead 2017
BRCA (BRCT domain), Shendure 2018
TPMT, Fowler 2018
GTPase HRas, Kuriyan 2017
PTEN, Fowler 2018
HIV env protein (BG505), Bloom 2018
Levoglucosan kinase, Whitehead 2015
HIV env protein (BF520), Bloom 2018
TIM Barrel, T. maritima, Matthews 2017
Small ubiquitin-related modifier 1, Roth 2017
HSP90, Bolon 2016
Translation initiation factor IF-1, Kishony 2016
Hepatitis C NS5A, Sun 2014
Ubiquitin, Bolon 2014
Ubiquitin, Bolon 2013
UBE4B (U-box domain), Klevit 2013
Influenza hemagglutinin, Bloom 2016
Flu polymerase PA subunit, Sun 2015
Thiamin pyrophosphokinase 1, Roth 2017
Calmodulin-1, Roth 2017
Levoglucosan kinase, Whitehead 2015
MAPK1/ERK2, Johannessen 2016
C
Spearman correlation
1.0
0.8
0.6
0.4
0.2
0.0
Experimental Pearson
Experimental Spearman
Autoregressive
DeepSequence
EVmutation
Independent
beta-lactamase, Ranganathan 2015
beta-glucosidase, Abate 2015
Kanamycin kinase APH(3')-II, Mikkelsen 2014
TIM Barrel, S. solfataricus, Matthews 2017
BRCA (Ring domain), Shendure 2018
SUMO-conjugating enzyme UBC9, Roth 2017
IGP dehydratase, Kondrashov 2018
BRCA (BRCT domain), Shendure 2018
TPMT, Fowler 2018
PTEN, Fowler 2018
HIV env protein (BG505), Bloom 2018
HIV env protein (BF520), Bloom 2018
HSP90, Bolon 2016
Ubiquitin, Bolon 2014
Ubiquitin, Bolon 2013
Influenza hemagglutinin, Bloom 2016
Ubiquitin, Fraiser 2016
Flu polymerase PA subunit, Sun 2015
MAPK1/ERK2, Johannessen 2016

**Supplementary Fig. 3.** Comparison of model performance across datasets. **A.** The Spearman correlation distributions of predictions compared to experimental measurements for each model. Two fragment lengths were used to build the HMM models (0.5 and 0.7), and only 0.5 is displayed in Figure 2. Two hidden sizes (24 and 48) were tested for SeqDesign; 48 was chosen for further study. Mean and quartile values are displayed for each model in the box-and-whisker plot. **B.** A comparison of model prediction Spearman correlations, including Envision LOPO prediction correlations. BRCA1 RING and UBE4B were validated but not tested by Envision due to poor validation performance. **C.** A comparison of model predictions' Spearman correlation with every dataset for which experimental Spearman or Pearson correlations are available. Pearson correlations have been included where a Spearman correlation is unavailable.

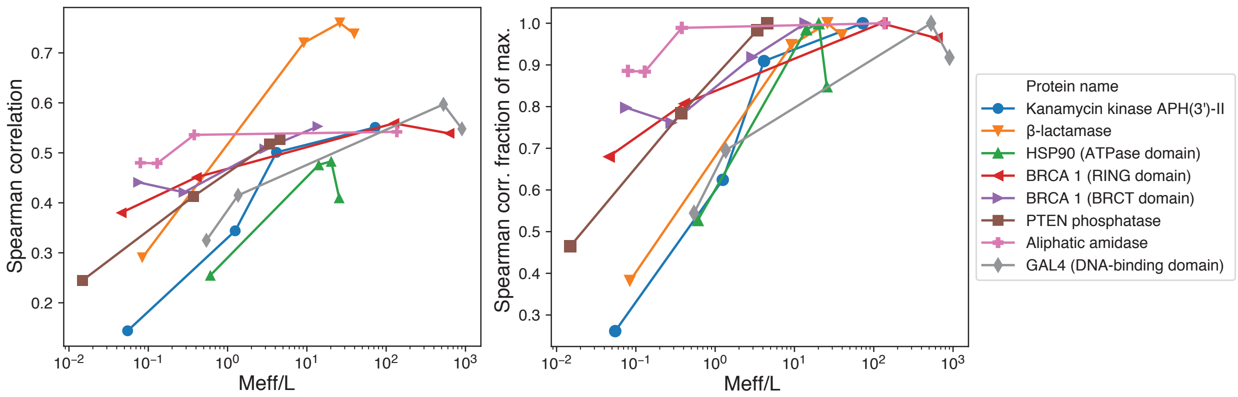


**Supplementary Fig. 4.** Spearman correlations for models trained on alignments of 8 protein families at 4 depths. Spearman correlations shown as-is (left) or normalized to the highest observed correlation for each family (right). Meff/L is the length-normalized number of nonredundant sequences after weighting sequences at 80% identity. Prediction accuracy for aliphatic amidase is nearly identical between a 151,555 sequence set (Meff=36,020; Meff/L=136; ρ=0.542) and a 3,982 sequence set (Meff=123; Meff/L=0.38; ρ=0.536).

SeqDesign:

Zabetakis, et al (2013)

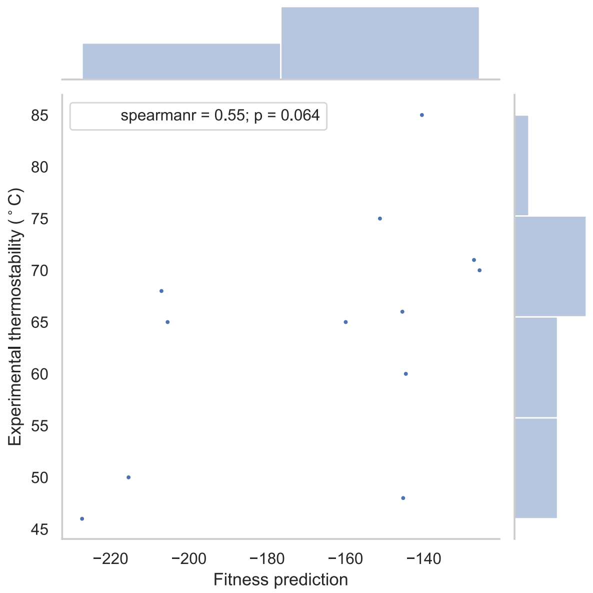


Kunz, et al (2017)

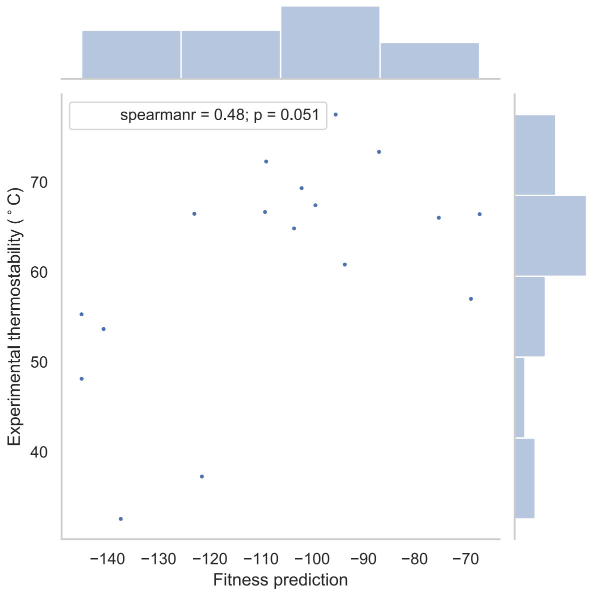


Shriver-Lake, et al (2017)

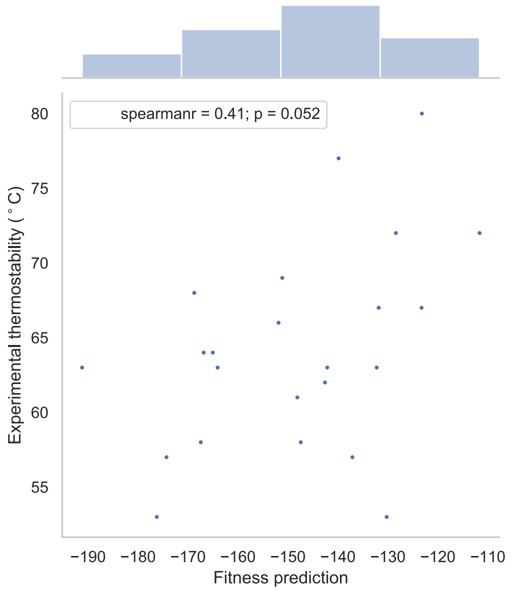


Turner, et al (2015)

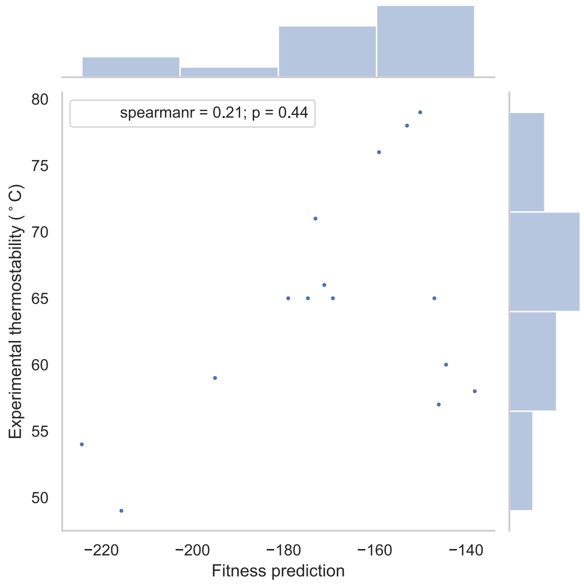

HMM:

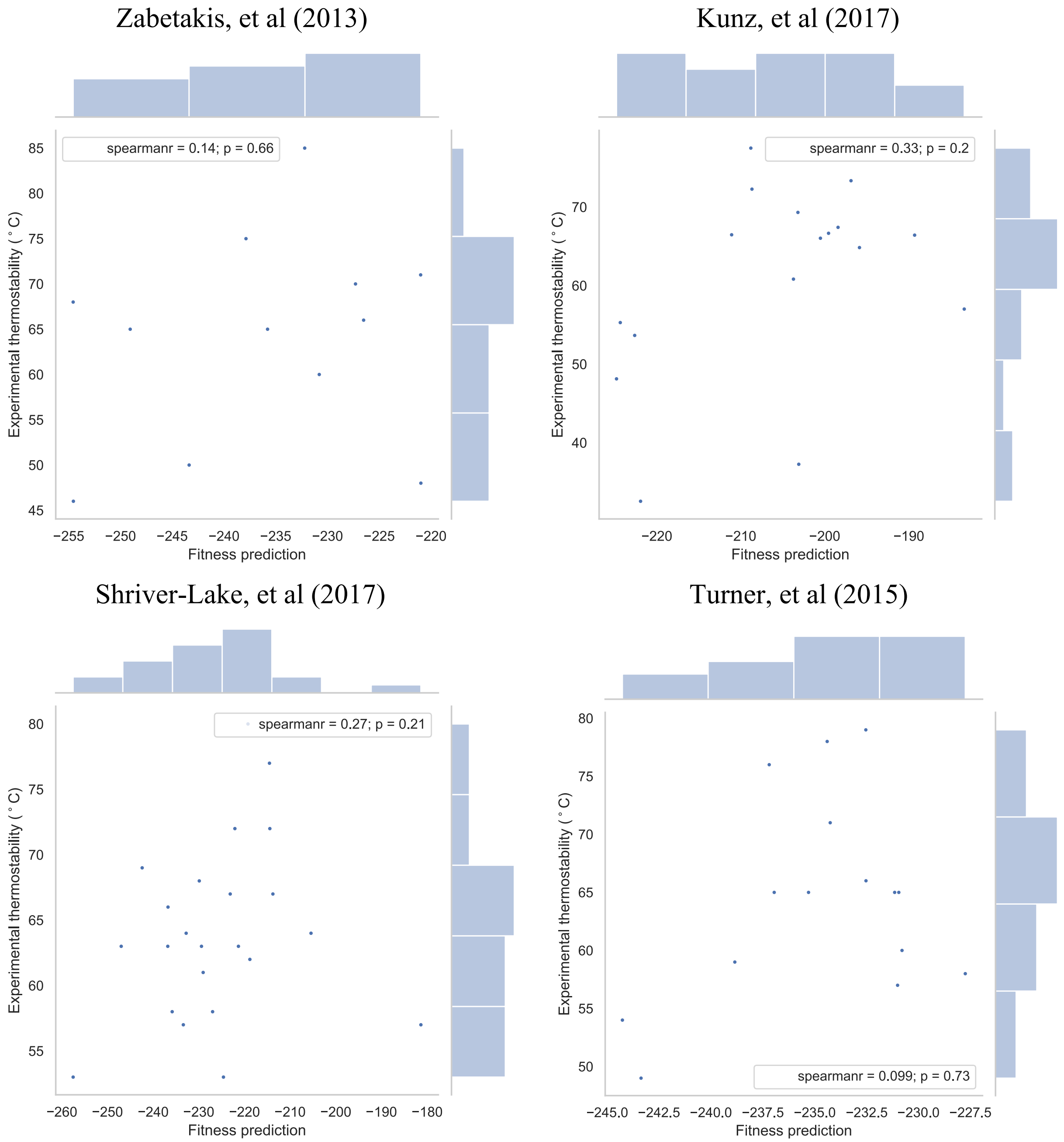


**Supplementary Fig. 5.** Fitness prediction of the SeqDesign and HMM models vs. experimental thermostability measurements from nanobody thermostability datasets for which there are at least ten data points. Thermostability measurements are in degrees Celsius and the fitness predictions are reported as log probability scores of each nanobody sequence.

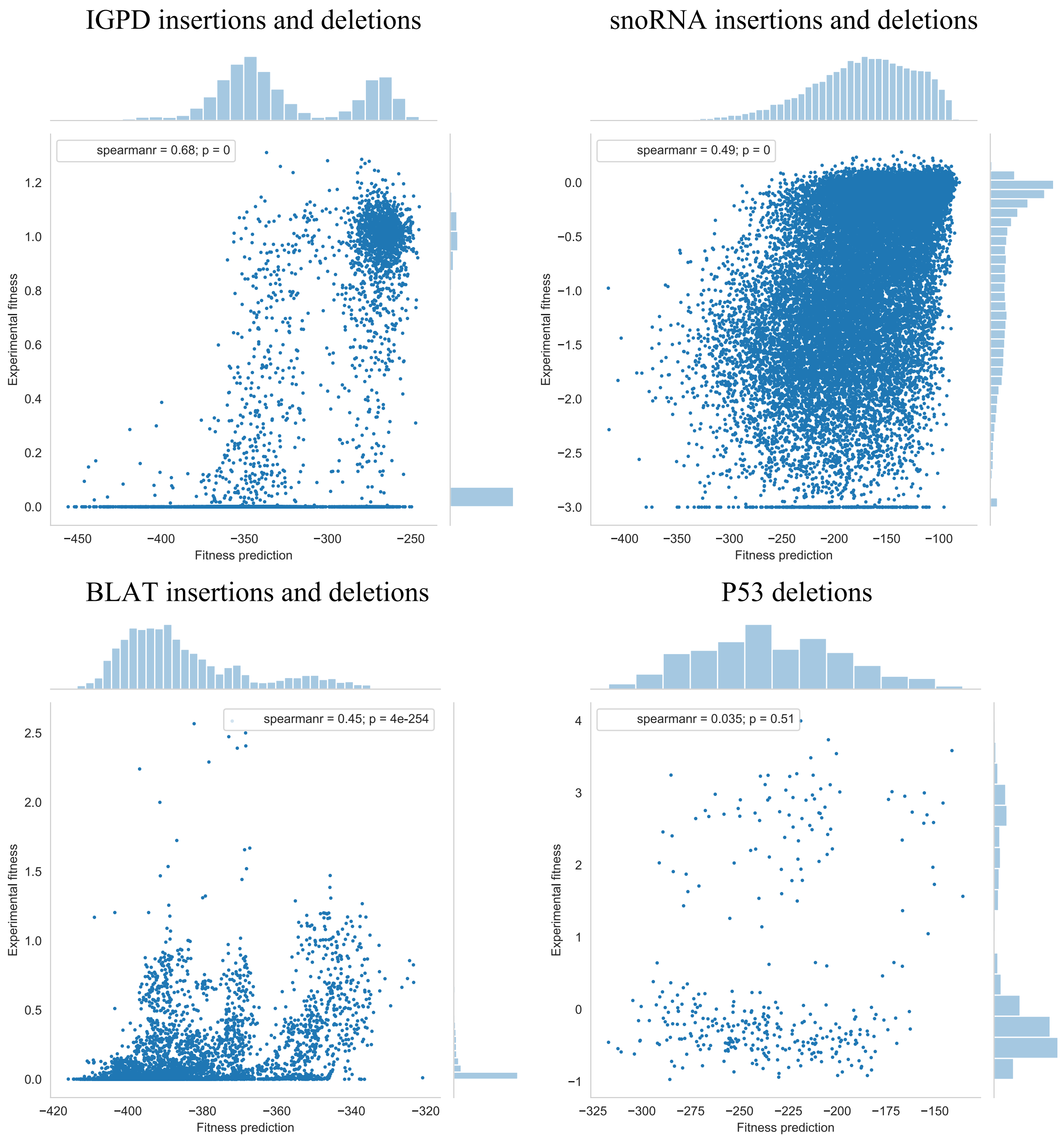


**Supplementary Fig. 6.** Indel mutation scan measurement comparisons for three proteins and one RNA: IGP dehydratase, snoRNA, $\beta$-lactamase and P53.

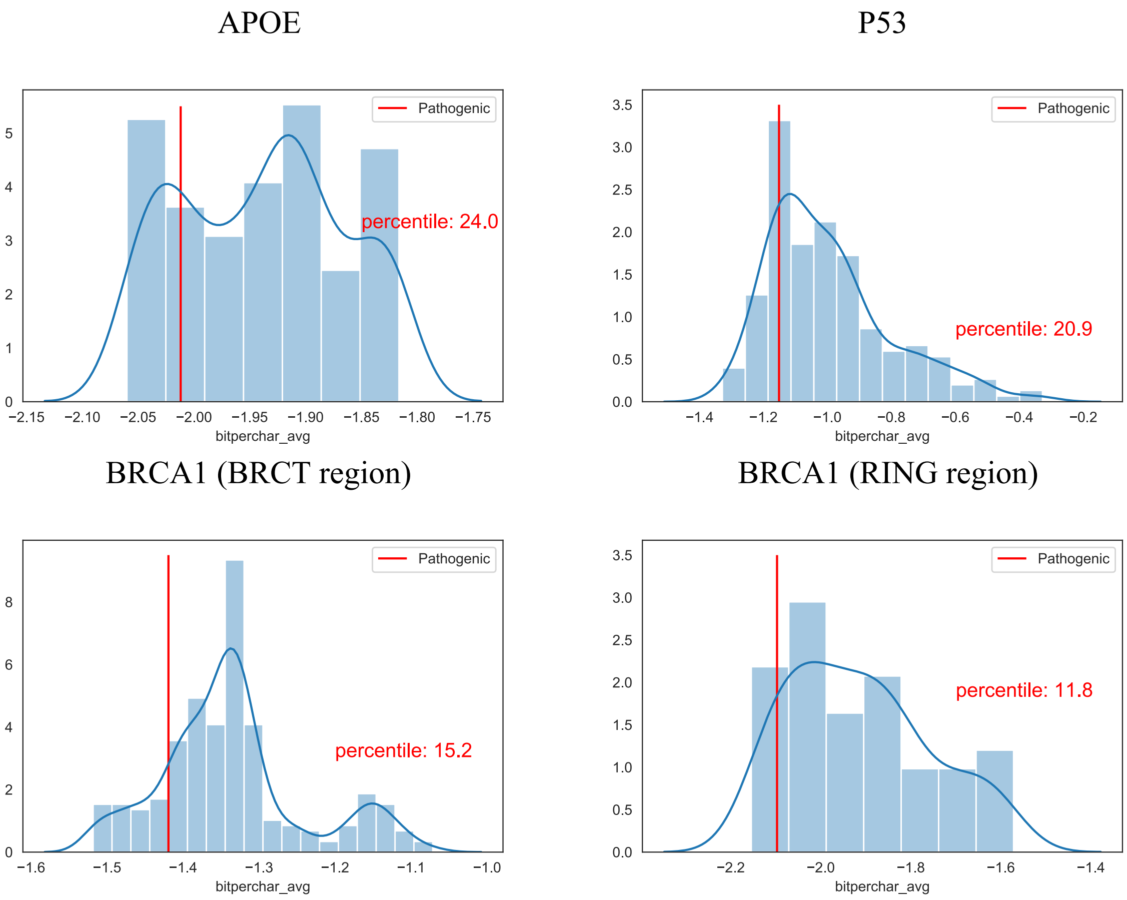


**Supplementary Fig. 7.** Pathogenic single amino acid deletions (as annotated by Clinvar) are predicted to be on the more deleterious spectrum of all possible single amino acid deletion effect predictions in a gene indicated in Alzheimer's (APOE), and two genes indicated in cancer (P53, BRCA1). Other single amino acid deletions that are predicted to be highly deleterious by the autoregressive model may be interesting to test for pathogenicity.

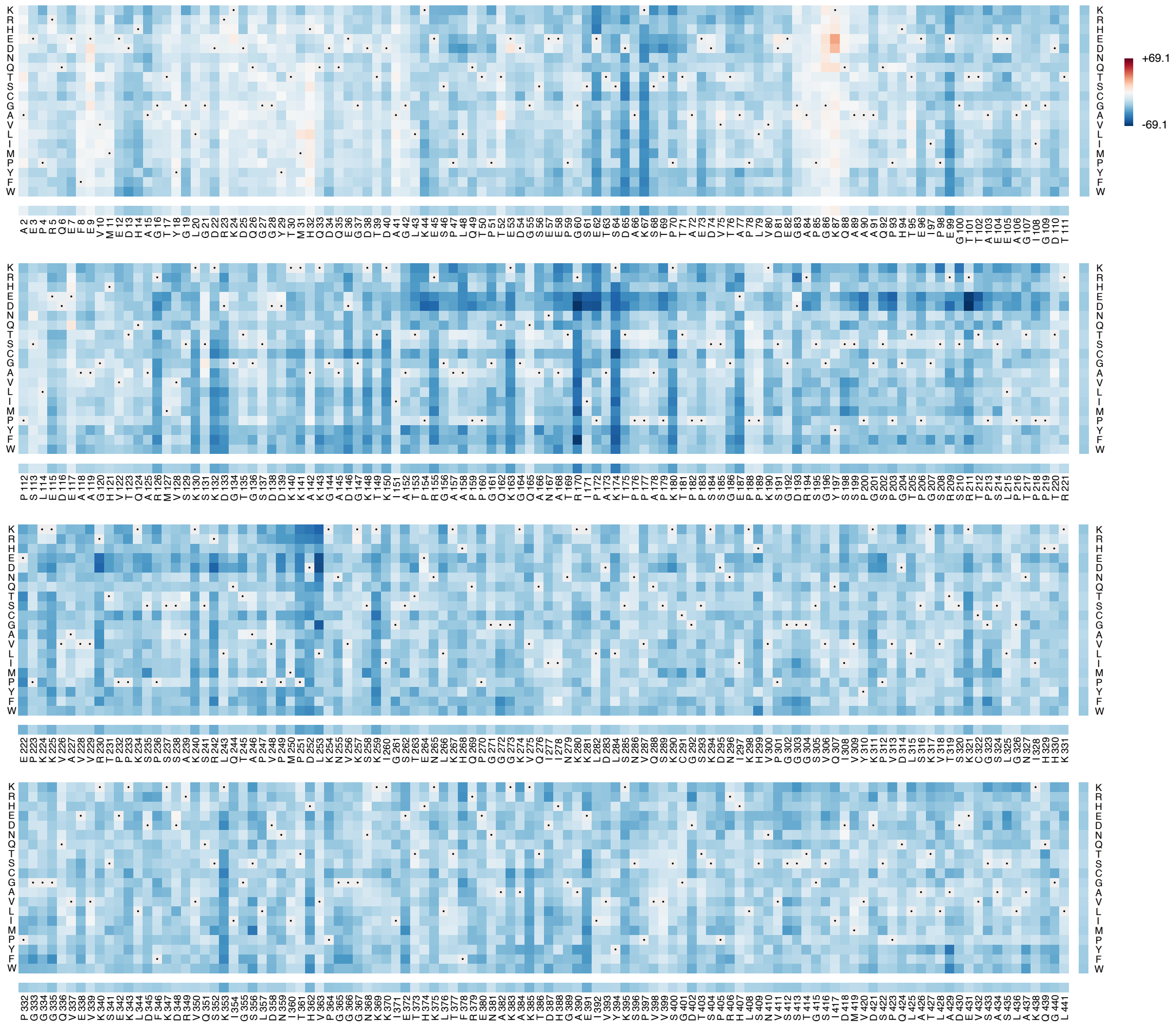
+69.1
-69.1

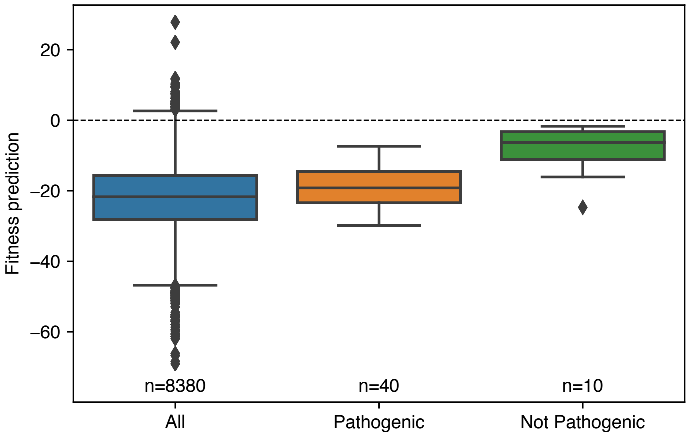
Fitness prediction
20
0
−20
−40
−60
n=8380
n=40
n=10
All
Pathogenic
Not Pathogenic

**Supplementary Fig. 8.** (top) *In silico* mutation scan of all single mutants for the human Tau protein, isoform Tau-4 (P10636-8). (bottom) Distribution of fitness predictions relative to wild-type for all mutations and for known variants annotated as pathogenic or not pathogenic in the Alzforum repository (https://www.alzforum.org/mutations/mapt). Fitness predictions distinguish pathogenic and not pathogenic groups (two-tailed independent $t=-4.5$, $P=3.8\times10^{-5}$; AUC=0.60)

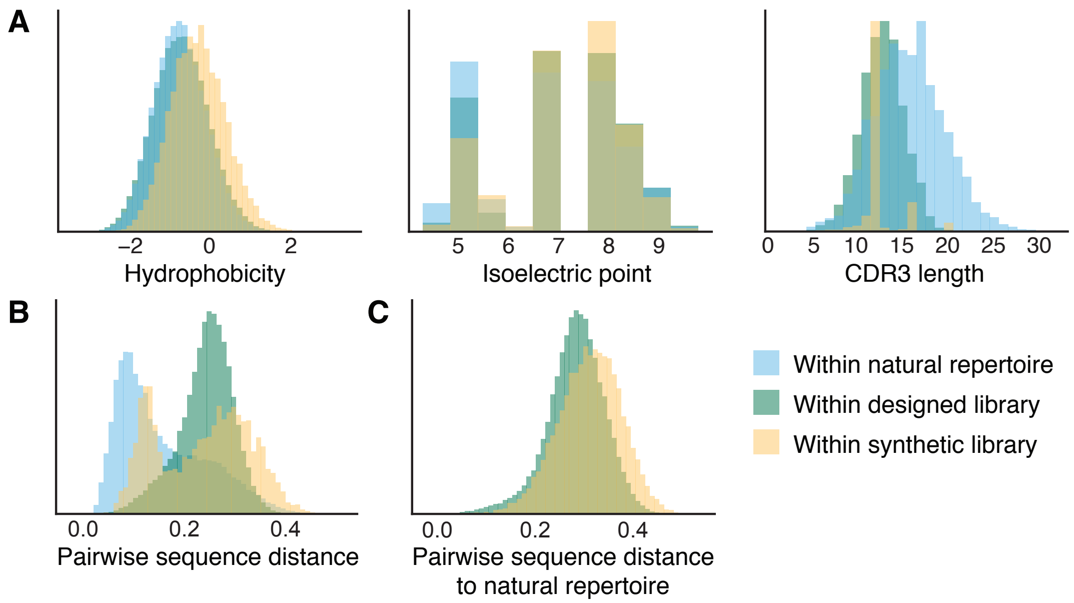


**Supplementary Fig. 9.** Compared to a synthetic library generated by codon randomization (McMahon et al., 2018), the designed library has more similar properties to the natural repertoire, while maintaining nearly the diversity of the synthetic library. a. Synthetic CDR3s have slightly different distributions of hydrophobicity and isoelectric point from the natural repertoire, and the synthetic library contains CDR3s of 3 lengths rather than a broad length distribution. b, c. The designed library contains nearly as much diversity as the synthetic library as measured by b) distances to the nearest neighbors within each library, and c) distances to the nearest neighbors in the natural llama single-domain antibody repertoire.

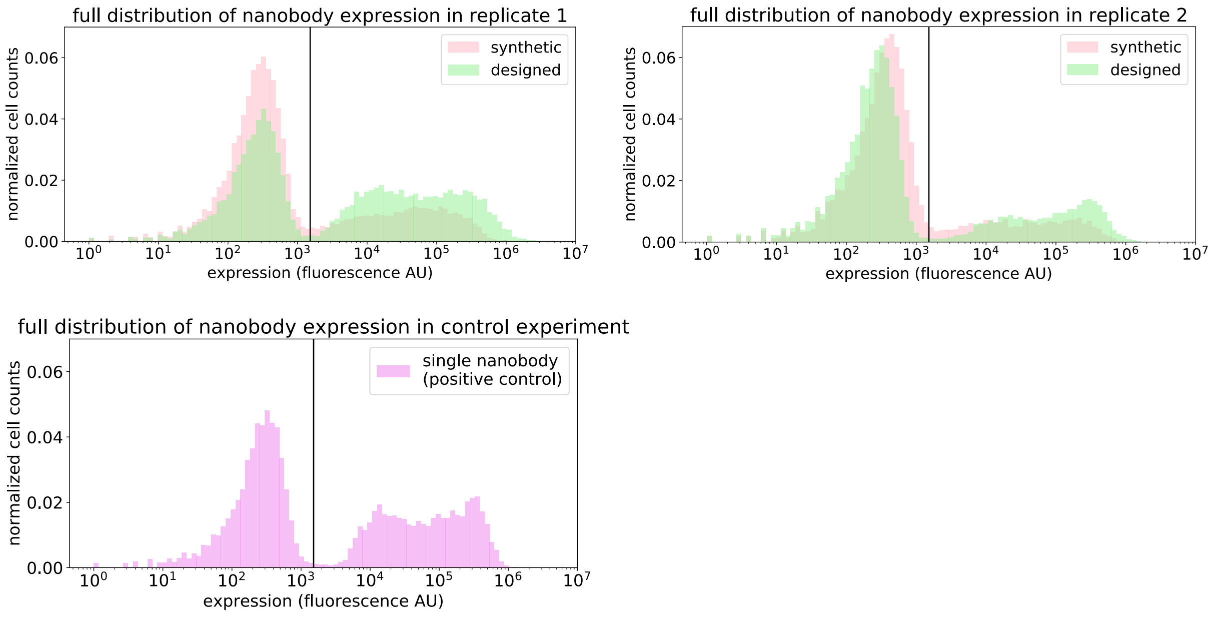


**Supplementary Fig. 10.** Full distributions of nanobody expression in the original synthetic library and our designed library. The designed library has a larger fraction of cells expressing nanobodies compared to the synthetic library (large difference in replicate 1 and small in replicate 2) and is closer to resembling the positive control. The vertical lines are the local minima between the non-expressing cells (the mode to the left of the line) and the expressing cells (to the right of the line). The distribution of the expressing cells are displayed in the main text in **Fig. 4a**.

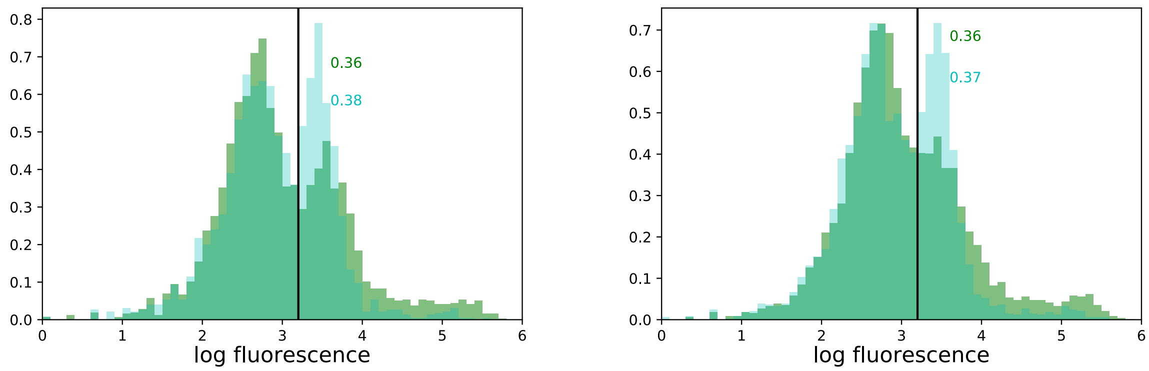


**Supplementary Fig. 11.** Experimental measurements of polyreactivity in the designed library (green) as compared to the original synthetic library (blue) shows similar, if not slightly lower proportions of poly-reactive nanobodies in the designed library when sorted for binding to a non-specific insect cell membrane reagent. The log experimental fluorescent measurements are shown on the x-axis, in two replicates (left and right).

SeqDesign nanobody library

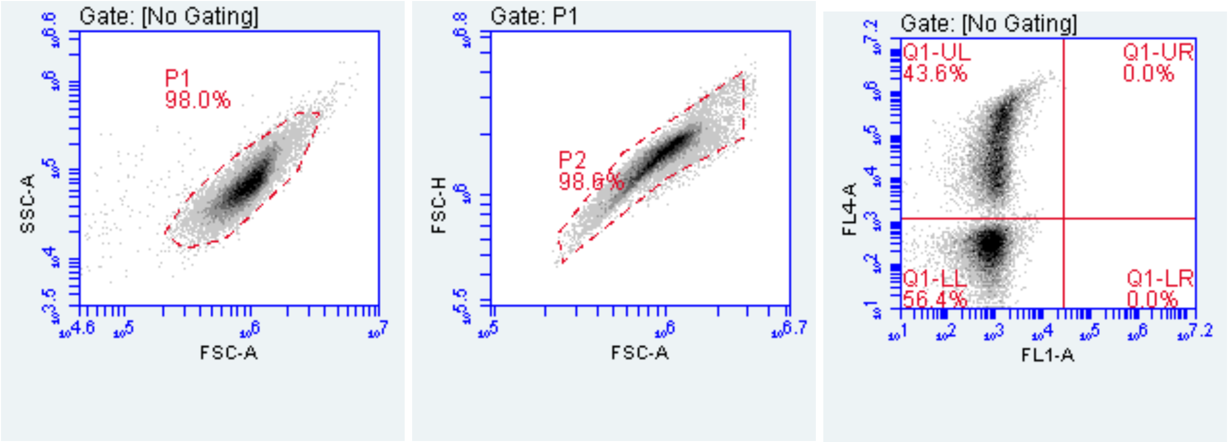


Synthetic nanobody library (McMahon, et al, 2018)

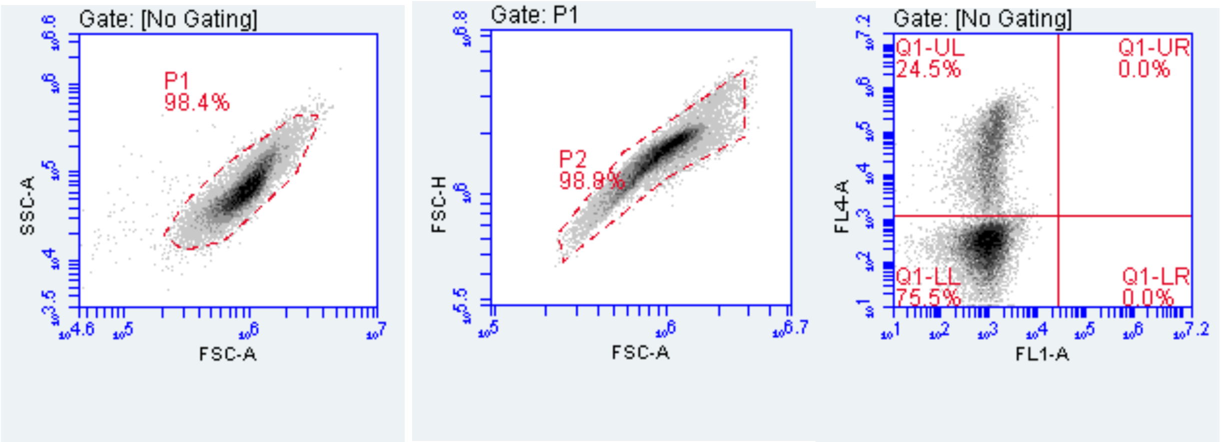


**Supplementary Fig. 12.** Example of flow cytometer yeast gating. Standard gating was used. Yeast form a single population in an FSC/SSC plot and were gated accordingly. Cells were further gated on an FSC-A/ FSC-H plot to exclude doublets. FL1-A uses AlexaFluor 647 and FL4-A uses AlexaFluor 488.

### Supplementary Tables

| Protein name | Uniprot ID (Nov 2015) | Measurement | # Mutations (mutant type) | Coverage | Number of replicates | Replicate correlations | Correlation notes | Reference |
|---|---|---|---|---|---|---|---|---|
| β-glucosidase | (Sequence from dataset) | Enzyme function | 3000 (single) | 1-501 | 2 | 0.97 | Pearson R, from paper | Romero et al., PNAS, 2015 |
| β-lactamase | BLAT_ECOLX | Growth | 3000 (single) | 24-286 | 1 | n/a | they report an estimated error per measureme nt | Firnberg et al., Mol Biol Evol, 2014 |
| β-lactamase | BLAT_ECOLX | Growth | 4997 (single) | 26-290 | 2 | 0.91 | Pearson R^2, from paper | Stiffler et al., Cell, 2015 |
| β-lactamase | BLAT_ECOLX | MIC | 990 (single) | 24-286 | 1 | n/a | Two experiment al conditions reported, 1 replicate each | Jacquier et al., PNAS 2013 |
| β-lactamase | BLAT_ECOLX | Growth | 4998 (single) | 24-286 | 1 | n/a | One replicate provided | Deng et al., JMB, 2012 |
| PSD95 (PDZ domain) | DLG4_RAT | Peptide binding | 1578 (single) | 311-393 | 1 | n/a | One replicate provided | McLaughlin et al., Nature, 2012 |
| GAL4 (DNA-binding domain) | GAL4_YEAST | Growth | 1196 (single) | 2-65 | 1 | n/a | One replicate provided | Kitzman et al., Nat Methods, 2015 |
| Influenza hemagglutinin | (Sequence from paper) | Viral replication | 10717 (single) | 2-565 | 3 | 0.66, 0.59, 0.59 | Pearson R^2, from paper | Doud & Bloom, Viruses, 2016 |
| HSP90 (ATPase domain) | HSP82_YEAST | Growth | 4324 (single) | 2-231 | 2 | 0.92 | Pearson R^2, from paper, but only for 20 amino acid positions | Mishra et al., Cell Reports, 2016 |
| Kanamycin kinase APH(3')-II | KKA2_KLEPN | Growth | 4582-4996 (single) | 1-264 | 2 | 0.91 | Pearson R, from paper | Melnikov et al., NAR, 2014 |
| DNA methylase HaeIII | (stabilized sequence based on MTH3_HAEAE) | Growth | 1957 (single, full), 1778 (single, filtered) | 2-330 | 1 | n/a | One replicate provided | Rockah-Shmuel et al., PLOS Comp Bio, 2015 |
| Influenza polymerase PA subunit | (Sequence from paper) | Viral replication | 1822 (single) | 8-716 | 2 | 0.96 | Pearson R, from paper | Wu et al., PLOS Genetics, 2015 |
| Poly(A)-binding protein (RRM domain) | PABP_YEAST | Growth | 1188 (single), 36522 (double) | 126-200 | 1 | n/a | One replicate provided | Melamed et al., RNA, 2013 |
| Hepatitis C NS5A | POLG_HCVJF | Viral replication | 1632 (single) | 1994-2097 | 1 | n/a | One replicate provided | Qi et al., PLOS Pathogens, 2014 |
| Ubiquitin | RL401_YEAST | Growth | 1196 (single) | 2-76 | 2 | 0.93 | Pearson R^2, from paper | Roscoe et al, JMB, 2013 |
| Ubiquitin | RL401_YEAST | E1 reactivity | 1366 (single) | 2-76 | 2 | 0.96 | Pearson R^2, from paper | Roscoe et al, JMB, 2014 |
| UBE4B (U-box domain) | UBE4B_MOUSE | Ligase activity | 900 (single) | 1072-1173 | 1 | n/a | One replicate provided | Starita et al., PNAS, 2013 |

| | | | | | | | | |
|---|---|---|---|---|---|---|---|---|
| YAP1 (WW domain 1) | YAP1_HUMAN | Peptide binding | 363 (single) | 170-203 | 1 | n/a | One replicate provided | Araya et al., PNAS, 2012 |
| Aliphatic amidase | AMIE_PSEAE | Enzyme function | 4507-4554 (single) | 1-341 | 1 | 0.889 | Pearson R, from paper | Wrenbeck et al., Nat Commun, 2017 |
| TIM Barrell (T. thermophilus) | P84126_THETH | Growth | 1520 (single) | 44-238 | 1 | n/a | One replicate provided | Chan et al., Nat Commun, 2017 |
| TIM Barrell (S. solfataricus) | TRPC_SULSO | Growth | 1520 (single) | 44-235 | 2 | 0.946 | Pearson R, from paper, but only for 20 amino acid positions | Chan et al., Nat Commun, 2017 |
| TIM Barrell (T. maritima) | TRPC_THEMA | Growth | 1520 (single) | 40-230 | 1 | n/a | One replicate provided | Chan et al., Nat Commun, 2017 |
| Translation initiation factor IF-1 | IF1_ECOLI | Growth | 1274 (single) | 1-72 | 1 | n/a | One replicate provided | Kelsic et al., Cell Systems, 2016 |
| Mitogen-activated protein kinase 1 | MK01_HUMAN | Growth | 5463 (single) | 2-360 | 4-6 | 0.98 | Pearson R, calculated from raw data | Brenan et al., Cell Rep, 2016 |
| Hras | RASH_HUMAN | Enzyme function | 3040 (single) | 2-166 | 1 | n/a | One replicate provided | Bandaru et al., Elife, 2017 |
| Ubiquitin | RL401_YEAST | Growth | 1142-1201 (single) | 2-76 | 2 | 0.79 | Pearson R^2, from paper | Mavor et al., Elife, 2016 |
| BRCA 1 (RING domain) | BRCA1_HUMAN | Growth | 492 (single) | 1631-1855 | 2 | 0.88 | Spearman R, from paper | Findlay et al., Nature 2018 |
| BRCA 1 (BRCT domain) | BRCA1_HUMAN | Growth | 1185 (single) | 1-101 | 2 | 0.88 | Spearman R, from paper | Findlay et al., Nature 2018 |
| Thiopurine S-methyltransferase | TPMT_HUMAN | Protein stability | 2659 (single) | 1-245 | 8 | 0.67 | Spearman R, from paper | Matreyek et al., Nat Gen, 2019 |
| Phosphatidylinositol 3,4,5-trisphosphate 3-phosphatase and dual-specificity protein phosphatase PTEN | PTEN_HUMAN | Protein stability | 3014 (single) | 2-403 | 8 | 0.62 | Spearman R, from paper | Matreyek et al., Nat Gen, 2019 |
| Levoglucosan kinase (stabilized) | (Stabilized sequence based on B3VI55_LIPST) | Growth | 6327 (single) | 1-439 | 2 | n/a | Plot provided, but no correlation metric | Klesmith et al., ACS Synth Bio, 2015 |
| Levoglucosan kinase | B3VI55_LIPST | Growth | 6541 (single) | 1-439 | 2 | n/a | Plot provided, but no correlation metric | Klesmith et al., ACS Synth Bio, 2015 |
| HIV env protein (BF520) | (Sequence from paper) | Viral replication | 12236 (single) | 30-691 | 3 | 0.59,0.60,0.64 | Pearson R, from paper | Haddox et al. Elife, 2018 |
| HIV env protein (BG505) | (Sequence from paper) | Viral replication | 12217 (single) | 30-699 | 3 | 0.76,0.77,0.78 | Pearson R, from paper | Haddox et al. Elife, 2018 |
| Imidazoleglycerol-phosphate dehydratase (His3) | HIS7_YEAST | Growth | 496137 (1-28 mutations) | 1-220 | 2 | 0.9 | Pearson R^2, from paper | Pokusaeva et al., PLOS Genetics, 2019 (bioRxiv 2017) |
| Calmodulin-1 | CALM1_HUMAN | Yeast growth | 1730 (single) | 2-149 | 1 | n/a | One replicate provided | Weile et al., MSB, 2017 |

| | | | | | | | | |
|---|---|---|---|---|---|---|---|---|
| Thiamin pyrophosphokinase 1 | TPK1_HUMAN | Yeast growth | 2608 (single) | 2-243 | 1 | n/a | One replicate provided | Weile et al., MSB, 2017 |
| Small ubiquitin-related modifier 1 | SUMO1_HUMAN | Yeast growth | 1329 (single) | 2-101 | 1 | n/a | One replicate provided | Weile et al., MSB, 2017 |
| SUMO-conjugating enzyme UBC9 | UBC9_HUMAN | Yeast growth | 2281 (single) | 2-158 | 2 | 0.78 | Pearson R, from paper | Weile et al., MSB, 2017 |

**Supplementary Table 1.** Deep mutational scans included in the paper and information regarding the experimental data, including sequence information, number of mutations, and experimental replicate measurements.

| Protein name | Uniprot ID | Original # sequences | Original Meff | Bitscore | # sequences | Meff | Sequence coverage | Meff/L | Original spearman | Spearman correlation |
|---|---|---|---|---|---|---|---|---|---|---|
| Kanamycin kinase APH(3')-II | KKA2_KLEPN | 29705 | 9380 | 0.3 | 56823 | 17439 | 240 | 72.66 | 0.554 | 0.551 |
| | | | | 0.5 | 4314 | 1034 | 250 | 4.14 | | 0.501 |
| | | | | 1 | 1636 | 318 | 255 | 1.25 | | 0.344 |
| | | | | 1.5 | 119 | 14 | 255 | 0.05 | | 0.144 |
| β-lactamase | BLAT_ECOLX | 14691 | 3667 | 0.3 | 40391 | 10013 | 253 | 39.58 | 0.741 | 0.738 |
| | | | | 0.5 | 27648 | 6691 | 257 | 26.04 | | 0.760 |
| | | | | 1 | 14614 | 2346 | 257 | 9.13 | | 0.720 |
| | | | | 1.5 | 1539 | 22 | 262 | 0.08 | | 0.290 |
| HSP90 (ATPase domain) | HSP82_YEAST | 23260 | 2586 | 0.3 | 45433 | 5672 | 222 | 25.55 | 0.473 | 0.410 |
| | | | | 0.5 | 43521 | 4517 | 223 | 20.26 | | 0.483 |
| | | | | 1 | 32922 | 3176 | 225 | 14.12 | | 0.476 |
| | | | | 1.5 | 4719 | 137 | 226 | 0.61 | | 0.255 |
| BRCA 1 (RING domain) | BRCA1_HUMAN | 39129 | 6304 | 0.3 | 251665 | 37215 | 60 | 620.25 | 0.571 | 0.539 |
| | | | | 0.5 | 78851 | 9508 | 76 | 125.11 | | 0.559 |
| | | | | 1 | 1125 | 44 | 109 | 0.40 | | 0.451 |
| | | | | 1.5 | 845 | 5 | 110 | 0.05 | | 0.380 |
| BRCA 1 (BRCT domain) | BRCA1_HUMAN | 8331 | 1883 | 0.3 | 13773 | 2829 | 205 | 13.80 | 0.545 | 0.553 |
| | | | | 0.5 | 3272 | 635 | 215 | 2.95 | | 0.508 |
| | | | | 1 | 1099 | 63 | 223 | 0.28 | | 0.421 |
| | | | | 1.5 | 808 | 17 | 225 | 0.08 | | 0.441 |
| PTEN phosphatase | PTEN_HUMAN | 8487 | 994 | 0.3 | 15397 | 1475 | 323 | 4.57 | 0.536 | 0.527 |
| | | | | 0.5 | 13168 | 1100 | 325 | 3.38 | | 0.518 |
| | | | | 1 | 1314 | 146 | 392 | 0.37 | | 0.413 |
| | | | | 1.5 | 709 | 6 | 403 | 0.01 | | 0.245 |
| Aliphatic amidase | AMIE_PSEAE | 76187 | 19554 | 0.3 | 151555 | 36020 | 264 | 136.44 | 0.578 | 0.542 |
| | | | | 0.5 | 3982 | 123 | 324 | 0.38 | | 0.536 |
| | | | | 1 | 2145 | 44 | 340 | 0.13 | | 0.479 |
| | | | | 1.5 | 2087 | 27 | 340 | 0.08 | | 0.480 |
| GAL4 (DNA-binding domain) | GAL4_YEAST | 22980 | 7026 | 0.3 | 156727 | 42406 | 47 | 902.26 | 0.622 | 0.548 |
| | | | | 0.5 | 98386 | 25868 | 49 | 527.92 | | 0.597 |
| | | | | 1 | 354 | 91 | 67 | 1.36 | | 0.415 |
| | | | | 1.5 | 101 | 40 | 74 | 0.54 | | 0.325 |

**Supplementary Table 2.** Spearman correlations of predictions with experiments using training sequence sets derived from alignments at four different bitscores. Meff is the number of nonredundant sequences after weighting sequences at 80% identity. Sequence coverage is the length of the focus sequence covered by the alignment, including non-focus columns that would have been excluded by alignment-based models.

| Reference | Dataset description | Number of datapoints |
|---|---|---|
| Zabetakis, et al., PLoS One, 2013 | Different regions stitched together | 12 |
| Shriver-Lake, et al., Toxicon, 2017 | Various point mutations introduced in different frameworks | 23 |
| Turner, et al., Biotechnol Rep, 2015 | Variety of mutation types (15 without disulfide linkage) | 19 |
| Kunz, et al., Biochim Biophys, 2017 | Potentially stabilizing variants designed based on sequence analysis | 17 |

**Supplementary Table 3.** Description of thermostability datasets for llama nanobody sequences used to validate the model's predictive capacity for nanobodies.

| Protein name | Uniprot ID | # Mutations | Replicate correlations | Correlation notes | Reference |
|---|---|---|---|---|---|
| PTEN | PTEN_HUMAN | 340 (deletions) | 0.58 | | Mighell et al., Am. J. Hum. Genet., 2018 |
| HIS3 | HIS7_YEAST | 5711 (deletions), 391 (insertions) | n/a | 197 isolated strains show r = 0.82 for fitness as measured by competition vs growth rate | Pokusaeva et al., PLOS Genetics, 2019 |
| snoRNA | CL00100 (RFAM) | 3896 (insertions), 22772 (deletions), 33144 (missense) | 0.87 | many different measurements with different correlations, but the two with the same conditions: small, 30, glu has a correlation of 0.87 | Puchta et al., Science, 2016 |
| P53 | P53_HUMAN | 357 (deletions), 5858 (missense) | n/a | n/a | Kotler et al., Mol Cell, 2018 |
| BLAT | BLAT_ECOLX | 262 (deletions), 4422 (insertions) | n/a | n/a | Gonzalez et al., J Mol Biol, 2019 |

**Supplementary Table 4.** Indel mutation scans that were used for validating the model's predictive power for sequences of different lengths. Experimental scans are included in the paper and their summary statistics.

| Step | Number of sequences (millions) |
|---|---|
| Generate sequences | 33.0 |
| Filter for reference final beta strand | 23.6 |
| Remove duplicate sequences | 21.7 |
| Remove duplicates of training set | 6.25 |
| Remove glycosylation motifs | 6.11 |
| Remove asparagine deamination motifs | 6.04 |
| Remove sulfur-containing amino acids | 3.69 |

**Supplementary Table 5.** Number of unique nanobody sequences remaining after each step of filtering of the designed nanobody library.

**Extended Data 1. (separate file)**
PTEN phosphatase deletion predictions with the autoregressive model trained on the PTEN sequence family.

**Extended Data 2. (separate file)**
IGP dehydratase insertion and deletion predictions with the autoregressive model trained on the IGP sequence family.

**Extended Data 3. (separate file)**
snoRNA insertion and deletion predictions with the autoregressive model trained on the snoRNA clan from RFAM.

**Extended Data 4. (separate file)**
Nanobody thermostability prediction measurements with the autoregressive model trained on naïve llama nanobody sequences.

**Extended Data 5. (separate file)**
In silico mutation scan of the Tau protein using the autoregressive model to generate predictive scores for all single missense mutations, including the disordered repeat region of Tau, which is indicated in neurodegenerative disease.

**Extended Data 6. (separate file)**
Predicted scores for Tau mutations categorized as pathogenic or not pathogenic in the Alzforum database. Mutations with uncertain significance were not included.
